# Modern microbial mats from the Chihuahuan Desert provide insights into ecological stability throughout Earth’s history

**DOI:** 10.1101/2021.11.18.469043

**Authors:** David Madrigal-Trejo, Jazmín Sánchez-Pérez, Laura Espinosa-Asuar, Valeria Souza

## Abstract

Microbial mats are complex ecological assemblages that are found in the Precambrian fossil record and in extant extreme environments. Hence, these structures are regarded as highly stable ecosystems. In this work, we assess the ecological stability in a modern, fluctuating, hypersaline pond from the Cuatro Ciénegas Basin. From the 2016 to 2019 metagenomic sampling of this site, we found that this microbial site is sensitive to disturbances, which leads to high taxonomic replacement. Additionally, the mats have shown to be functionally stable throughout time, and could be differentiated between dry and rainy seasonal states. We speculate that this microbial system could represent modern analogs of ancient microbial mats where functions were preserved over time, whereas composition was subject to diversification in the face of local and planetary perturbations.

## 1 Introduction

There is little to no doubt that life emerged early in Earth’s history, as suggested by geochemical signatures, biomarkers, microfossils and sedimentary structures from the early Archean (Lepot, 2020). Particularly, phototrophic microbial mats, alongside stromatolites, have been extensively present in the Archean rock record, as shown in the fossil evidence from the Dresser formation (3.48 Ga) (Noffke, Christian, et al., 2013), the Buck Reef Chert (3.42 Ga) (Tice and Lowe, 2004; Tice, 2009), and the Moodies group (3.22 Ga) (Noffke, Eriksson, et al., 2006; Homann, Heubeck, et al., 2015; Homann, Sansjofre, et al., 2018). Microbial mats are also found in modern environments; they are benthic, stratified, and self-sustaining biological communities of thousands of phylogenetically diverse microorganisms embedded in a matrix of extracellular polymeric substances (EPS) (Prieto-Barajas et al., 2018). Therefore, it is straightforward to infer that microbial mats have been thriving on Earth for more than ~3.5 Ga, in spite of every threat posed to life.

Indeed, Earth’s history has been marked with gradual transitions and punctuated events that certainly disturbed the early biosphere. The early Sun, although 30% fainter than today, emitted high-frequency radiation, coronal mass ejections and solar cosmic rays by 2-3 orders of magnitude greater than present values (Obridko et al., 2020); geomagnetic polarity transitions would increase the solar wind and cosmic rays flux (Erdmann et al., 2021); asteroid impacts of bolides of 20-70 km in diameter struck Earth at 3.47–3.23 Ga and possibly until 3.0 Ga, way after the Late Heavy Bombardment (Lowe et al., 2014; Davatzes et al., 2019); surface chemistry shifted from a reduced state towards an oxidized world during the Grate Oxidation Event (2.43-2.22 Ga) (Gumsley, Chamberlain, et al., 2017; Poulton et al., 2021); global glaciation events were triggered by changes in the carbon cycle and solar heating (Tajika, 2007; Arnscheidt and Rothman, 2020); and large igneous provinces flooded the surface with effusive volcanism towards the end of the Archean and during the Phanerozoic (Mole et al., 2018; Gumsley, Stamsnijder, et al., 2020). Each of these environmental pressures could potentially erradicate life from Earth, yet, life (as we know it) survived.

The success of microbial mats and stromatolites as biological structures can only be understood in terms of ecological stability; namely, the community response to disturbances, which could be dissected into the degree to which a community is insensitive to perturbations (ecological resistance) and the rate at which a community restores to the pre-disturbed state (ecological resilience) (Shade et al., 2012; Song et al., 2015). Environmental disturbances can be classified into pulses and presses if the perturbation is a discrete, short-term event, or a continuous, long-term transition, respectively (Bender et al., 1984; Shade et al., 2012). Microbial community stability is a topic of interest for a wide array of systems and disturbances, such as dry-rewetting events (Kolda et al., 2019), differences in water level (Ren et al., 2019), temperature variations (García-García et al., 2019; Okonkwo et al., 2020), chemical stress (Jiang et al., 2020), shifting redox patterns (Pett-Ridge and Firestone, 2005), and changes in salinity (Berga et al., 2017). Nonetheless, microbial community stability under the scope of early life geobiology is rarely explored.

In this work, we study the microbial system denominated as the Archean Domes, Cuatro Ciénegas, Mexico (Fig. 1). This pond is subject to extreme conditions, such as prolonged droughts, intense solar radiation, and major shifts in salinity and pH. Hypersaline microbial mats are among the best studied type of mats, and they have been widely recognized as analogs to the Archean Earth and, plausibly, early Mars (Wong, Smith, et al., 2015; Perl and Baxter, 2020; Saona et al., 2020). Hence, we took a metagenomic, uniformitarian approach to assess ecological stability and community dynamics from a three-year sampling to speculate the underlying processes and mechanisms that enable microbial communities to cope with multiple disturbances throughout Earth’s history.

**Fig. 1:**
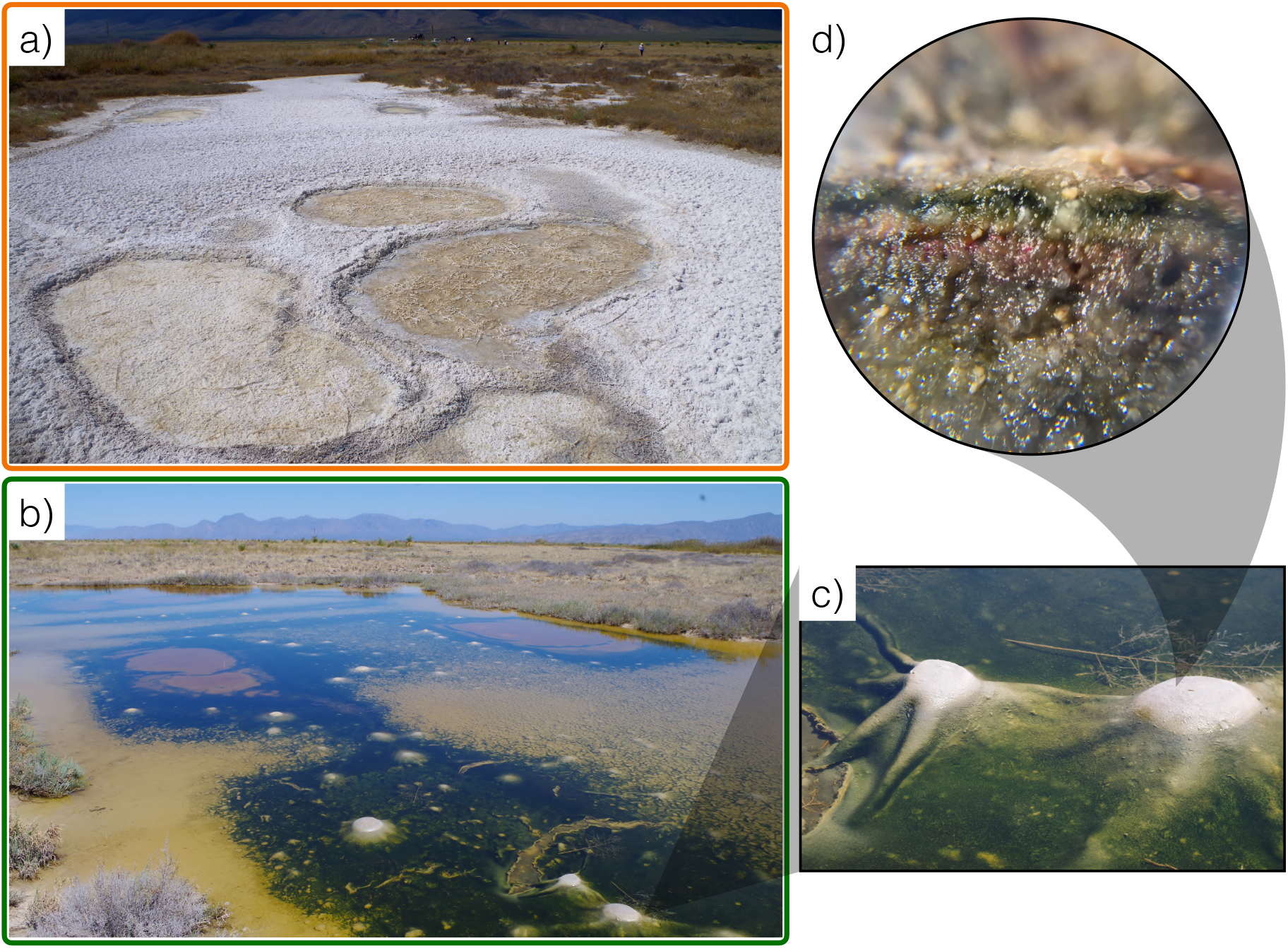
The Archean Domes microbial system. The pond displays different features during *a)* dry season (sampling of March 2019) and *b)* rainy season (sampling of March 2016). *c)* Detail of the dome structures. *d)* 10X magnification of a microbial mat; functional stratification and sediment grains can be appreciated at this scale.

## 2 Materials and methods

### 2.1 Study site and sample collection

The Archean Domes (26º49’41.7”N, 102º01’28.7”W) is a seasonal, water-fluctuating pond in Rancho Pozas Azules from Pronatura located at the eastern side of the Cuatro Ciénegas Basin, Coahuila, México (Site overview in Fig. S1: Supplementary material). This site was discovered in 2016, and was firstly described by Medina-Chávez et al. (2019) [unpublished], and Espinosa-Asuar et al. (2021) [unpublished]. During the rainy season, mostly during the months of August to September, the pond fills with water up to ~20 cm. Green mats emerge over the soil surface, building dome-like sedimentary structures up to 10-15 cm in diameter (Fig. 1b,c). From November to July, water evaporates and salt precipitation covers the pond completely, burying the microbial mats (Fig. 1a). Salinity is variable between the two states, transitioning from 52.5 PSU (as measured in the rainy season of 2016) when filled with water to salt saturation during the dry season. From a recent sampling in September 2021, we observed that green mats and gas filled structures start to quickly develop after the day of rainfall. Inside the domes and mats there are variable concentrations of methane (2.6-19.6 μg/L on the rainy season of 2016, 102-402 *μ*g/L on the dry season of 2017) and carbon dioxide (1.08-1.40 on the dry season of 2017). During dry season, pH is ~7, while on rainy season the pH rises to ~8.5-9.5 with the dissolution of salts.

We collected six samples of mats and associated sediment across a three-year period. During this time span, we got to collect three samples of each seasonal state: dry and rainy season. The mats from the dry season are from the sampling of April 2016, February 2017 and March 2019 (hereinafter denoted as M1_D16, M3_D17 and M5_D19, respectively). Mats from the rainy season are from the sampling of October 2016, October 2018 and September 2019 (hereinafter denoted as M2_R16, M4_R18 and M6_R19, respectively). Rainy season samples come directly from developing domes, whereas dry season samples derive from soil regions where mat was visible to the naked eye. As the rainy season is heavily contingent on the cyclone dynamics of the Gulf of Mexico, samples were taken at different times, ~1-2 weeks after a heavy rainfall to ensure a high level of water in the pond. To prevent contamination, samples were collected with gloves, sterile forceps and sterile conical tubes (50 mL) and stored in liquid nitrogen for their preservation prior to extract the DNA. Weather parameters were taken from the National Meteorological Service, CONAGUA, at the EMA station No. 15DBB372 in Cuatro Ciénegas (27º0’7.2”N, 102º4’22.7”W). Weather data is provided in Fig. S2 and Table S1: Supplementary material.

### 2.2 DNA purification and sequencing

From each sample, only the mat layer (~1 cm) was taken for DNA extraction. As the samples of the dry seasons contain a thick layer of salt, this layer had to be separated with a sterile scalpel to facilitate the extraction. We perform the extraction of total DNA from the six samples as reported in Purdy (2005). Purified DNA was sent to CINVESTAV-LANGEBIO for shotgun metagenomic sequencing. DNA libraries for Illumina paired-end sequencing were prepared for each sample without any amplification step. DNA from all samples was sequenced with Illumina MiSeq (2 x 300 base pair paired-end reads). The total number of paired-end reads per metagenome range from 4.7 to 28.0 Gbp per library and orientation (forward and reverse). Number of raw reads can be found in Table S2: Supplementary material.

### 2.3 Quality control, assembly and annotation of metagenomes

We preprocessed the raw reads with Trimmomatic v0.38 (Bolger et al., 2014) with a sliding window of 4, a Phred quality score of 30, minimum length of 35, and an average mean quality of 28. For each metagenome, reads were assembled into contigs to facilitate gene prediction. Forward and reverse paired reads, and individual forward and reverse with no pair, were assembled using MEGAHIT v1.1.1 (Li, Liu, et al., 2015) with minimum contig length of 500, k-min of 27 and k-step of 10 as suggested for highly-diverse metagenomes (Bandla et al., 2020; Yan et al., 2021). To control for sequencing depth bias, we used the minimum number of reads (1,288,875 reads) to sample the metagenomic datasets at random to normalize coverage for comparisons. Unassembled reads were collected with BBtools (Bushnell, 2020) and SAMtools v1.12 (Li, Handsaker, et al., 2009). For assembled contigs, gene prediction and subsequent taxonomic annotation was done with CAT v5.2 (Meijenfeldt et al., 2019). Additional information regarding quality control, metagenome assembly, processing of not assembled reads with MEGAHIT, and taxonomic annotation can be found in Table S2-S5: Supplementary material. CAT is a robust taxonomic annotator that integrates known software programs such as gene predictor Prodigal (Hyatt et al., 2010) and gene annotator DIAMOND (Buchfink et al., 2014) against the NCBI non-redundant database (NCBI Resource Coordinators, 2018) to give a deep gene taxonomic annotation. Since taxonomic annotation with CAT revolves against all kinds of predicted genes, we also used six ribosomal protein families (PF00177, PF00298, PF00573, PF00237, PF00163 and PF00318) to validate CAT results. We downloaded ribosomal genes’ seeds from Pfam database (Mistry et al., 2021). HMM profiles were built with HMMER v3.3 (Eddy, 2011), and hmmsearch was performed against all metagenomes (e-value 10^-6^). Ribosomal genes were annotated with DIAMOND, coupled with the NCBI non-redundant database. Overall functional profiling was done with SUPER-FOCUS (Silva et al., 2016) against the NCBI non-redundant database. We select resistance genes based on GO classification and download the amino acid sequences from UniProt database. Resistance query sequences were aligned with BLAST against all metagenomes. Finally, We selected key energy metabolisms and nutrient cycling as in Gutiérrez-Preciado et al. (2018). Protein families involved in each metabolic pathway were initially searched in UniProt (Bateman et al., 2021) and KEGG (Kanehisa et al., 2016) databases, and subsequently downloaded from Pfam. HMMER and BLAST (Altschul et al., 1990) analyses were performed for each protein family and for each metabolic pathway.

### 2.4 Normalization, statistical analyses and data visualization

We used R programming language (R Core Team, 2020) to run each statistical analysis, to normalize data and to generate figures. We list the libraries used as follows: ggplot2 v3.3.5 (Wickham, 2016) for overall plots, edgeR v3.34.1 for data normalization (Robinson et al., 2010), RAM v1.2.1.7 (Chen, Simpson, et al., 2018) for PCoA, PCA and CCA analyses, vegan v2.5-7 (Oksanen et al., 2020) for rarefaction curves and alpha-diversity metrics, UpSetR v1.4.0 (Lex et al., 2014) for upset plots, for differential expression analysis DESeq2 v1.32.0 (Love et al., 2014) and EnhancedVolcano v1.10.0 (Blighe et al., 2021), patchwork v1.1.1 (Pedersen, 2020) and fmsb v0.7.1 (Nakazawa, 2021) for radar charts, streamgraph v0.9.0 (Rudis, 2019) for streamgraphs, easyalluvial v0.3.0 (Koneswarakantha, 2021a) and parcats v0.0.3 (Koneswarakantha, 2021b) for alluvial plots, NetCoMi v1.0.2 for network analyses (Peschel et al., 2021), and umap v0.2.7.0 and dbscan v1.1-8 for clustering. Libraries BBmisc v1.11 (Bischl et al., 2017), dplyr v1.07 (Wickham et al., 2021), tidyr v1.1.4 (Wickham, 2021) were used for data manipulation. Gene abundances were normalized with the Relative Log Expression (RLE) method. PCoA and NMDS analyses for taxonomic groups were calculated with a Bray-Curtis measure. NetCoMi networks were built using SparCC measure, Bayesian-multiplicative replacement for zero handling and association threshold of 0.5. Phylum-level networks were built with the top 120 phyla, while genus-level networks were built with all the 250 core genera.

## 3 Results

### 3.1 Taxonomical characterization

We build rarefaction curves to evaluate diversity coverage for all samples. For genera richness, each sample reaches saturation and comparisons between them is suitable (Fig. 3S: Supplementary material). Open read-frames were predicted for reads, and further annotated for taxonomic classification with CAT. We detected 162 phyla, 2250 genera (across all samples), and more than 8,000 phylotypes per sample. Nevertheless, only 30-58% of the total predicted genes for each sample were classified, suggesting a considerable amount of potential novel taxonomic groups, which comprise the so called “microbial dark matter”. These potentially uncultured organisms have shown to be of importance in other hypersaline microbial mats (Wong, MacLeod, et al., 2020). Mean abundances per domain show consistent results between CAT and ribosomal gene annotation; for CAT taxonomic assignment we got mean abundances of: 85.24% for Bacteria, 14.43% for Archaea, and 0.3% for Eukaryota; whereas ribosomal gene annotation showed: 86.56% for Bacteria, 13.35% for Archaea, and 0.08% for Eukaryota.

Regarding the taxonomic composition, at the phylum level, samples consistently displayed Pro-teobacteria (23.51%), Euryarchaeota (11.42%), Bacteroidetes (10.26%), Firmicutes (4.35%), Cyanobacteria (3.30%), Spirochaetes (2.84%), Planctomycetes (1.99) and Chloroflexi (1.42) as the most abundant phyla (Fig. 2). The taxonomic annotation with ribosomal genes is also consistent with the phyla relative abundances of CAT annotation (taxonomic profile based on ribosomal proteins is shown in Figure S4: Supplementary material). Taxonomic profiles seem to vary between each sample; most noticeable, with the increase of Euryarchaeota for the 2019 samples. Overall, the Archean Domes have a high diversity as seen in Chao (143-271), Shannon (2.5-3.1) and inverse Simpson (4.6-8.3) indexes.

**Fig. 2:**
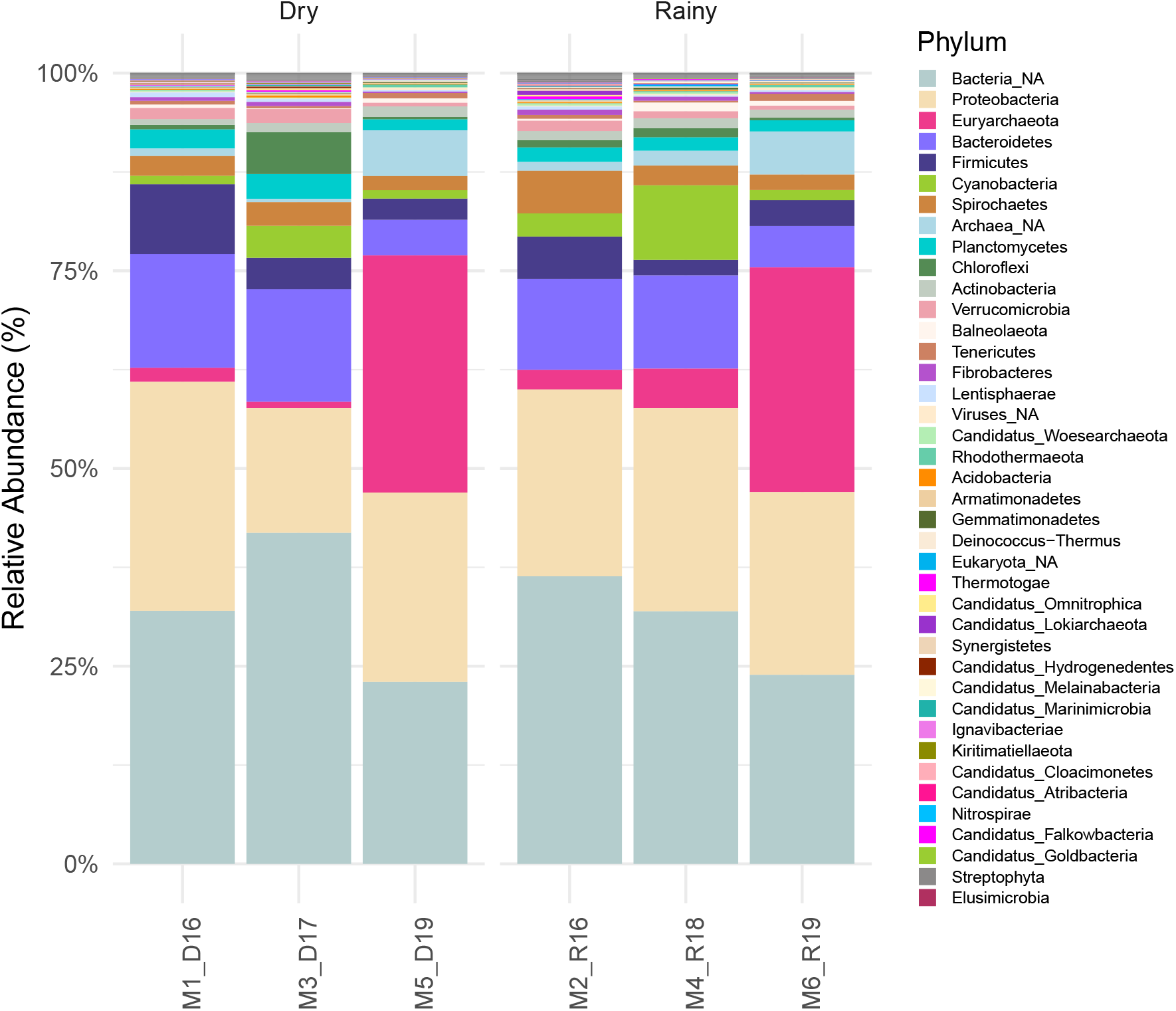
Taxonomic profile of the Archean Domes. Only the top abundant phyla are displayed. Not annotated phyla were grouped into NA category.

At the genus level, we find *Coleofasciculus* as the most abundant Cyanobacteria between all samples, which is widely known as a key mat-forming genus in sandy environments (Noffke, 2010; Ramos et al., 2017; Prieto-Barajas et al., 2018; Cardoso et al., 2019). Other cyanobacterial genera such as *Leptolyngbya*, *Halothece*, and *Phormidium* are also abundant between samples, and have also been previously reported in microbial mats (Ramos et al., 2017; Sohm et al., 2020; Brenes-Guillén et al., 2021). Anaerobic, halophilic, sulfate-reducing members of the Deltaproteobacteria such as *Desulfonatronovibrio*, *Desulfonatronospira* and *Desulfovermiculus* also appear in abundance in the Archean Domes samples.

(Kuever, 2014). Other relevant taxonomic genera present in the samples include *Halorubrum* (Euryarchaeota), *Halanaerobium* (Firmicutes), *Spirocheta* (Spirochetes), *Chitinispirillum* (Fibrobacteres), and *Tangfeifania* (Bacteroidetes). From the 2250 total genera found in the system, only between 16-19 for each sample belong to the abundant genera, that is, with an abundance >1%. In contrast, between 426-619 genera have abundances <0.1%, and belong to the so called “rare biosphere”. Rare taxa account for the 11.2-18.9% of the whole community, whereas abundant taxa comprises the 43.3-67.6%. Therefore, although taxa that are abundant only consists of a few genera, these taxa often build most of the microbial community biomass (Fig. S5: Supplementary material). Moderately abundant taxa (>0.1% and <1%) sits between the abundant and rare, with a relative abundance of 19.6-37.7% in the samples studied.

### 3.2 Functional characterization

Coding sequences were functionally classified in order to infer potential functions. As expected, basic functions shared between all living beings are widely distributed among all samples, such as carbohydrate (14.5%), amino acid (11.9%), protein (8.9%), DNA (5.9%), RNA (5.0%), and fatty acids and lipids (3.1%) metabolisms; Other processes regarding cofactor, vitamins, and pigments (10.7%), cell wall and capsule (4.2%), respiration (3.9%), and stress response (3.8%) are also among the top functions for all samples. Stress response genes in higher abundance might reflect that the community is subject to ceaseless environmental pressures (Varin et al., 2012; Le et al., 2016). Fig. 3a shows the difference in function abundance between samples for every major process according to SUPER-FOCUS classification. Overall, samples appear to be similar among them, despite some functions with differential distribution among the samples, such as amino acid, fatty acids and lipids, central, secondary, and RNA metabolisms.

**Fig. 3:**
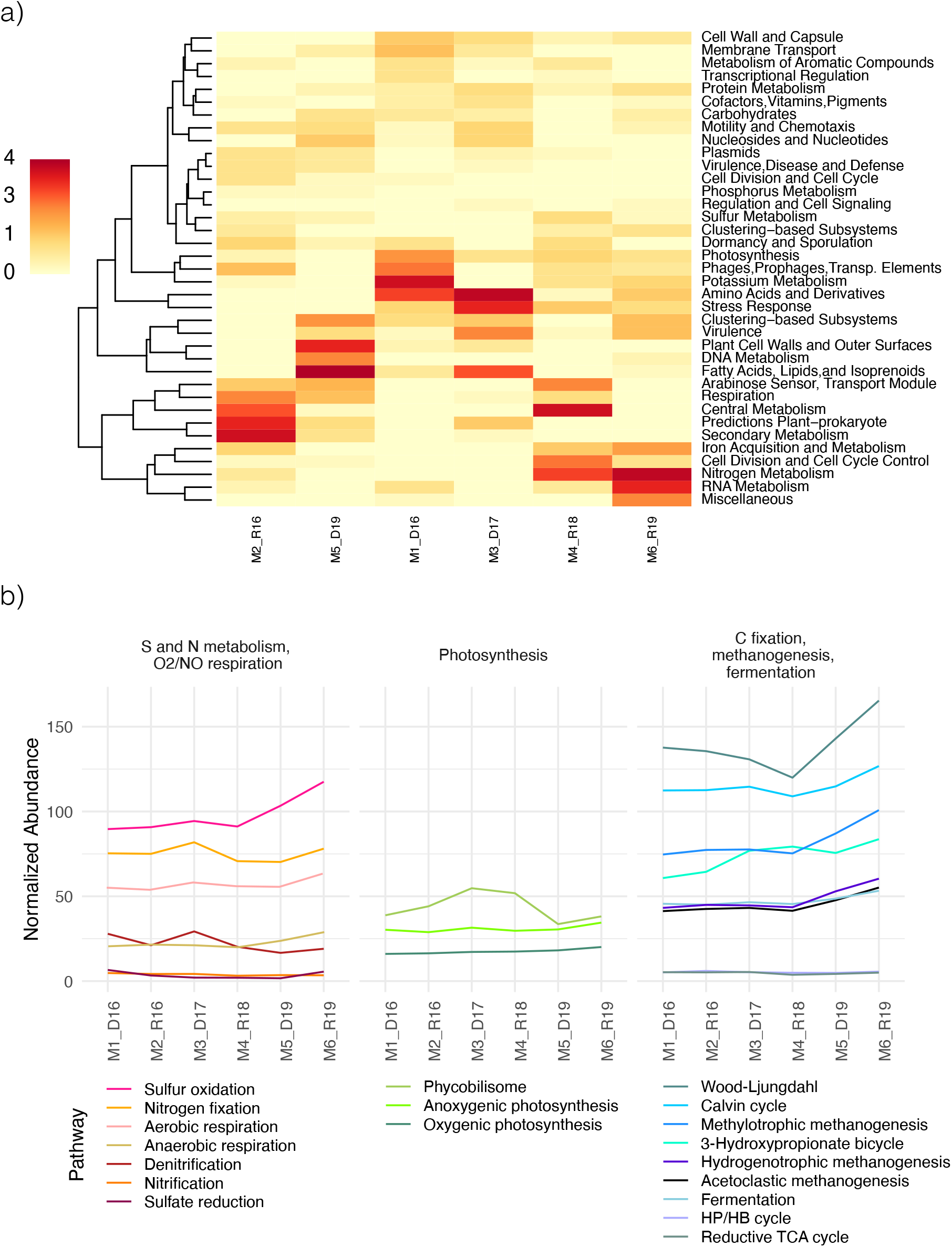
Potential functional profile based on metagenomic inference. *a)* Heatmap of SUPER-FOCUS major functions, with color intensity reflecting differences in abundance between samples. *b)* Normalized abundance of selected pathways based on pathway-specific Pfam protein groups. HP/HB=3-hydroxypropionate/4-hydroxybutyrate, TCA=tricarboxylic acid.

We inspect the function of the stress response genes present at the Archean Domes. Based on GO classification, we identify resistance genes related to pH, alkaline, acidic, salt, dormancy, and endosporulation conditions. Alkaline and salt resistance genes were the most abundant, with a mean proportion of 56.5% and 31.5%, respectively (Fig. S6: Supplementary material). This behavior is expected, since salt and pH fluctuate considerably between seasons, and might exert a selection pressure on the organisms thriving on this site.

Pfam protein groups were used to infer energy metabolisms and nutrient cycling within the mat samples. Based on normalized abundance, Wood-Ljungdahl pathway rules carbon metabolism among the mat, followed by the Calvin cycle. These results are consistent with other microbial mats previously described, and Wood-Ljungdahl dominance has been regarded as a result of energy limitation, since this mechanism of carbon fixation is inefficient compared to other pathways (Gutiérrez-Preciado et al., 2018; Wong, White, et al., 2018; Kurth et al., 2021). Anoxygenic photosynthesis genes dominate over those specific to oxygenic photosynthesis, while sulfur oxidation and nitrogen fixation are potentially the main processes for sulfur and nitrogen metabolisms. Dissimilatory sulfate reduction is portrayed as a process with low gene abundances, despite the highly abundant sulfate reducing bacteria previously described; as such, metabolism inference based on gene abundances should be taken cautiously. Further reconstruction of full pathways would provide more accuracy in the relative abundances.

### 3.3 Community dynamics through time and seasonal comparison

On account of the morphological changes of the pond in response to environmental perturbations, we conduct statistical analyses to evaluate if samples have higher resemblance to those collected in the same seasonal state. Chao, Shannon, and inverse Simpson indexes were calculated for each sample to evaluate alpha diversity, and no statistical significance was found between seasons (Wilcoxon Rank Sum test: Chao1 p=0.4, Shannon *p*=1, Inverse Simpson *p*=0.8). Moreover, principal coordinates analysis (PCoA) and non-metric multidimensional scaling (NMDS) at the genus and order level showed no seasonal aggregation of samples (Fig. S7: Supplementary material). Finally, Canonical correspondance analysis (CCA) was performed with the environmental variables provided by the EMA meteorological station (Fig. S8: Supplementary material). From this analysis, roughly, 2017-2018 samples were driven by precipitation, whereas 2019 samples were driven by wind speed and humidity. This could be non-conclusive due to the low sample number. Nonetheless, two groups seem to have formed: one less closer to each other from samples of 2016 - 2018, and other more closely arranged which comprises the samples from 2019. This result is expected, as taxonomic profiles from the dry and rainy seasons of 2019 showed similar compositions (Fig. 2), particularly, the increase in the Archaea relative abundances.

Co-occurrance networks were built to inspect general properties at the phylum level (Table S7 and Fig S9: Supplementary material). Both seasons mainly show two clusters which might be associated to groups of highly interacting organisms or functional guilds with niche overlapping to some degree. During the rainy season, several phyla from both groups transition to build a third cluster. Hence, it is possible that phyla interact differently between themselves depending on environmental conditions. Network metrics have been used to evaluate resilience and resistance within microbial communities. For instance, our networks shows a positive edge of 49.017 and 48.77 during the dry and rainy seasons, respectively. A high positive/negative ratio in microbial networks, such as those found in these networks, has been interpreted to aid in community stability, by avoiding feedback loops in taxa with overlapping niches (Hernandez et al., 2021). Furthermore, modularity has been considered as a measure of community stability, diminishing the propagation of perturbations through the network (Hernandez et al., 2021). The Archean Domes microbial mats seem to change in modularity between the dry season (0.01) and rainy season (0.07) states; lower modularity during the dry season might reflect the exposure and vulnerability of the system relative to when the mats are wet. The small sample size might induce spurious correlations in the microbial networks, and further samplings for the following seasons will support this analysis.

Seasonal patterns in community composition are not straightforward to follow, and evaluating the community dynamics through the years might provide a plausible underlying explanation for this. Fig. 4a show the community composition changes through the years. As briefly stated previously, one of the most noticeable changes through the years was a rise of Archaea (from 1-4% to 33%) in the samples of 2019. Consequently, Bacteria reduced their abundance up to ~65%, a third less from previous years. The virus followed the same tendency as the archaea druing 2019, in a subtle rise of abundance (0.08-0.2% to 0.4%). The Eukaryota had an apparent seasonal pattern in the first two years (2016-2018), continuing with a steady state in 2019. Since the increased abundance of Archaea was considerable, the dynamic between 2016-2018 is visually lost. Taking into account only the abundance shift between 2016-2018, all domains presented a possible seasonal pattern, where archaeas, eukaryotes and viruses rose proportionally in the rainy season compared to the dry one. To explore which organisms may drive these seasonal patterns, we examined phylum and genus proportion across time. Phyla with a prominent shift were Spirochaetes, Proteobacteria, Cyanobacteria, Cloroflexi, Bacteroidetes, and the Euryarchaeota. Euryarchaeota became one of the main abundant groups in the communities of 2019 (from 2% to 28%). In contrast, Cyanobacteria, Chloroflexi and Bacteroidetes showed a diminished abudnance during the same year. Spirochaetes had a rise in October 2016 to end in a constant frequency in the following samples.

**Fig. 4:**
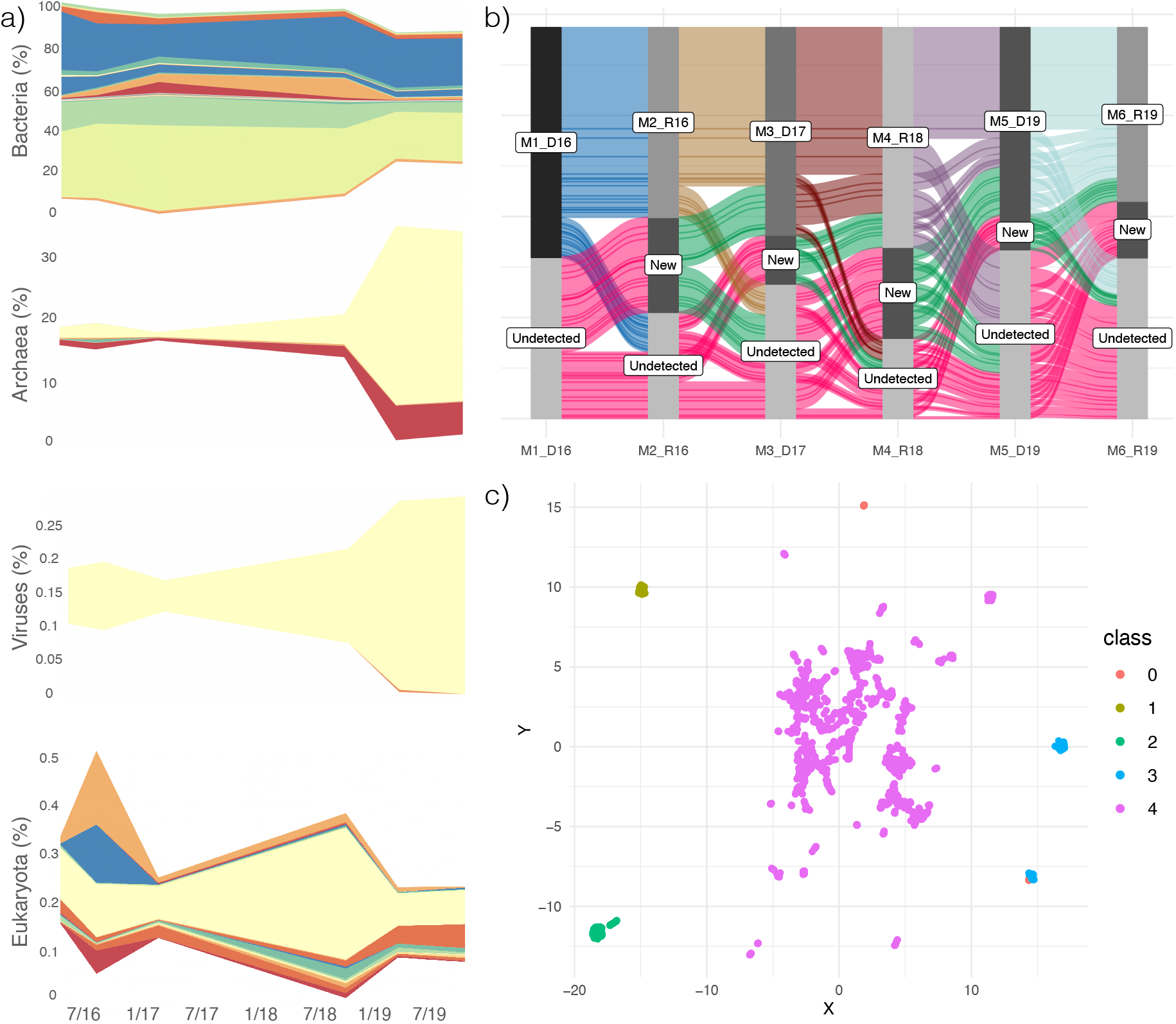
Taxonomic replacement and community dynamics throughout the years. *a)* Changes in superkingdom abundances from 2016 to 2019. Colors within each graph depict phyla. *b)* Community dynamics at the genus level. Flows show genera that remain, appeared or disappeared from the system through the samples. *c)* UMAP dimension reduction and HDBSCAN clustering technique applied on the differences in genus abundance from each sample.

At the genus level, the taxonomic replacement is even more noticeable (Fig. 4b). For each sampling, we could observe two main phenomena: i) some genera are present in every sample (core genera), while ii) most genera are new additions or become undetected between each sample. As a matter of fact, taxonomic replacement becomes increasingly complex with each new sample, which reflects how the community has changed since the first sampling in April 2016. We delve deeper into core dynamics in the following section. We were interested to evaluate which taxa are key in driving the community to new compositional states, based on the differential abundances between samples. Coupling UMAP, a nonlinear dimensionality reduction method, with HDBSCAN, Hierarchical Density-Based Spatial Clustering of Applications with Noise), we find groups that might be leading the community dynamics (Fig. 4c). First, the main cluster contains most of the genera, with the inclusion of all the abundant taxa (1734 genera in class 4). In contrast, four small groups with fewer genera in each one (22, 116, 200, and 182 genera in classes 0,1,2, and 3, respectively. Group composition is supplied as Supplementary material in a csv file). These groups are made up entirely of genera belonging to the rare biosphere, and shifts in their abundance seem to be major ecological drivers in the ecosystem. This result further support the relevance of the rare biosphere in microbial communities, as they could XXX (Jousset et al., 2017).

We assess if functional categories differentiate communities between seasons. Normalized abundances for general functions (system 1 level based on SUPER-FOCUS classification) showed that dry and rainy function abundance is essentially the same, with slight differences in abundance (Fig. 5a). This behavior is expected, as major functions, with fundamental roles in every microbe, are always present for the survival of the community. As described previously, most genes are associated to carbohydrate, amino acids, and protein, metabolisms as well as processes cofactors, vitamins, and pigments. Nevertheless, PCoA for these data with a Bray-Curtis measure do arrange them into seasonal groups, although ordination is sparse (Fig. 5b). Further sampling will support the predictability of this clustering method.

**Fig. 5:**
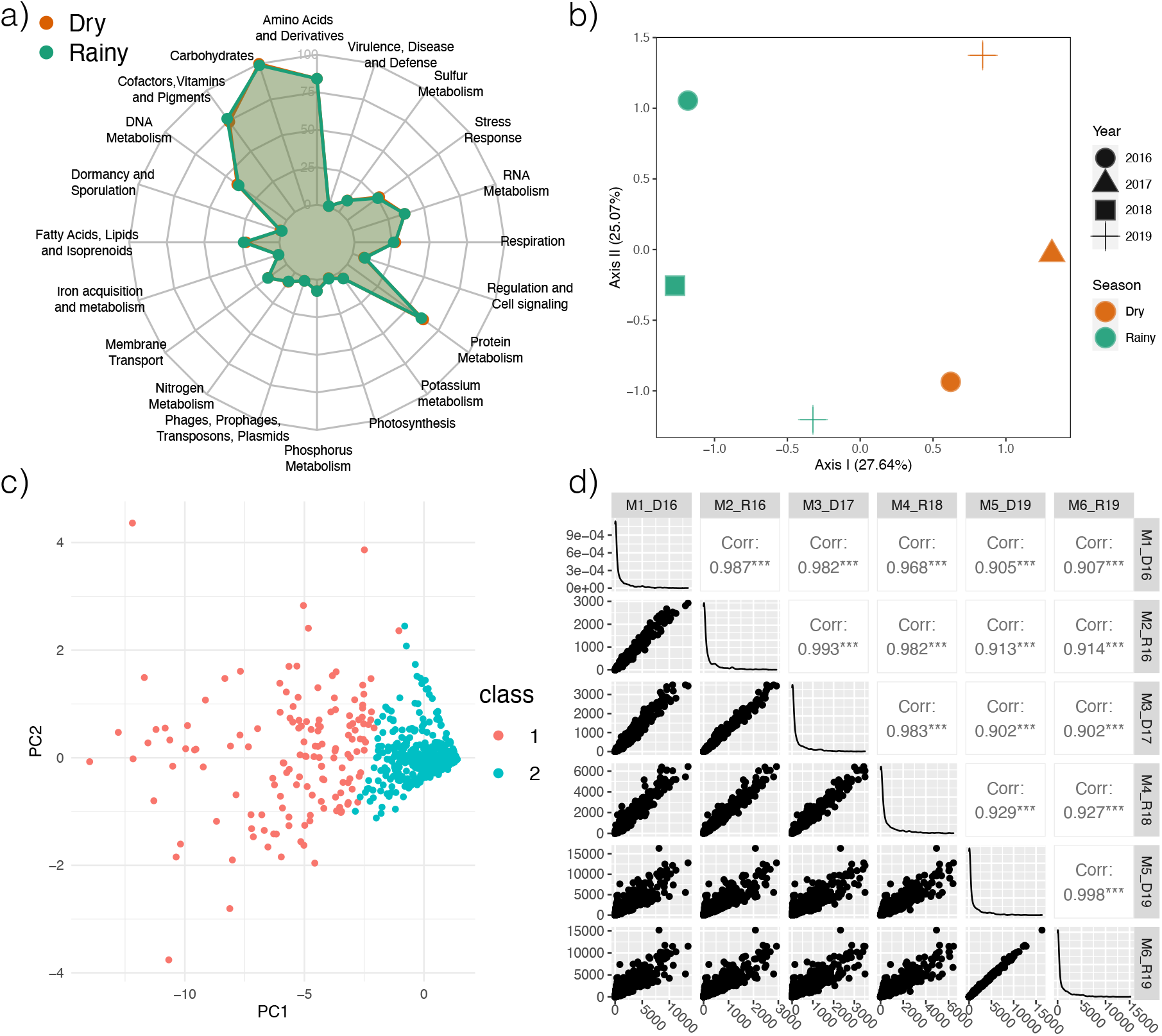
Functional comparison between the years and functional changes throughout the years. *a)* Dry and Rainy season comparison based on Top SUPER-FOCUS processes. *b)* PCoA with Bray-Curtis measure, where samples are grouped, to some extent, by season. *c)* PCA analysis showing main groups of functions by k-means clustering. *d)* Correlations of function abundance between each sample, where each sample is more similar to the adjacent ones in time.

We modify a differential expression analysis to adapt it to our metagenomic using the classification defined by SUPER-FOCUS. Although none of the metabolic subsystems had a significant difference between seasons (p > 0.5), there were some processes that had a higher or lower abundance as seen by their fold change (Log2 Fold Change>abs[2.5], Fig. S10: Supplementary material). In the dry season, there were three slightly more abundant functions: the pentose phosphate pathway of plants, the alpha-acetolactate operon, and the biotin biosynthesis. From the pentose phosphate pathway, we had the glucose 6 phosphate dehydrogenase, the key enzyme of the Oxidative Pentose Phosphate Pathway (OPPP), which is related to the response of short- or long-term exposure to drought stress in plants (Landi et al., 2016). The alpha-acetolactate operon has been described as a component in the mixed acid fermentation, done by some bacteria such as *Bacillus subtilis*, to produce acetoin in the absence of nitrate (Härtig and Jahn, 2012); this could be associated to a shortage of nutrients in the dry season. Lastly, biotin biosynthesis is an important process, since biotin is a key cofactor in the fatty acids and amino acid metabolisms, as well as in the replenishment of the tricarboxylic acid cycle (Salaemae et al., 2016). For the rainy season, some functions with a higher fold change were: the acyl homoserine lactone (AHL) inducer, which is involved in primary quorum sensing signals by Gram-negative bacteria (Parsek et al., 1999); the phage carbon metabolism Auxiliary metabolic genes (AMGs), which consist of phage strategies for resource management during host infection (Thompson et al., 2011; Warwick-Dugdale et al., 2019); some archaeal hydrogenases, involved in carbon fixation (Hedderich, 2004); prenylated indole alkaloids production from actinomycetes, which have multiple biological functions such as antifungal and antibacterial activity (Netz and Opatz, 2015); and lastly, chlorophyll degradation related genes. Some of these functions could be directly associated with the presence of the green, cyanobacterial built, layer seen in the rainy seasons, such as in the phage-cyanobacterial AMGs (Thompson et al., 2011), the quorum sensing for biofilm formation (Herrera and Echeverri, 2021), and the chlorophyll degradation.

Moreover, we were wondering how functions have changed over time and if there is a group of functions that leads global patterns in the community. According to k-means and hierarchical clustering, two main groups of functions were predicted. PCA analysis showed how these functions are projected, where high abundant functions are sparsely distributed in the plot, and most functions with low abundance functions were tightly clustered together (Fig. 5c). Each function’s class is provided in the Supplementary material. Comparing function abundance across samples suggest that functions are more similar between adjacent samples (Fig. 5d). In consequence, the correlation cloud appears to be scattering when samples are more distant in time. For instance, sample M1_D16 showed higher correlation with sample M2_R16 than with the last sample from 2019 (M6_R19). This result further suggest that functions are changing between samples, and that cumulative changes in functions differ drastically from the initial function state. As previously stated, further sampling may reinforce this hypothesis.

### 3.4 The core community

Taxonomic composition and functions change through time to some extent, as described in the previous section. Still, there is a core community shared between all samples and seasons. The (global) core community consists of 250 genera out of the 2250 total genera across the samples, just about ~11% of the total diversity found in the Archean Domes (Fig. 6a). These genera can be portrayed as microbes with high physiological plasticity, able to cope with both dry and rainy season environmental conditions (Pett-Ridge and Firestone, 2005). Seasonal cores were identified, that is, genera that only appeared in rainy or in dry season exclusively. Unlike the core community, seasonal cores were particularly small, with just 1 and 10 genera for dry and rainy seasons, accordingly. Every genus in the seasonal cores have a low abundance (<0.01%), and belong to the rare biosphere during each season. The organisms found only in rainy samples comprise several Alphaproteobacteria (*Croceicoccus*, *Shimia*, *Rhodoplanes*, and *Polymorphum*), Gammaproteobacteria (*Teredinibacter* and *Allochromatium*), Bacteroidetes (*Ohtaek-wangia*), Cyanobacteria (*Geminocystis*), one Euryarchaeota (*Methanosalsum*) and a novel genus of Planctomycetes (Candidatus *Jettenia*) previously described in an anammox bioreactor (Mardanov et al., 2019). Among the genera present only in rainy season, it is noticeable the presence of the phototrophs *Allochromatium* (purple sulfur bacteria), *Rhodoplanes* (photoheterotroph) and *Geminocystis* (Cyanobacteria) (Imhoff, 2014; Marcondes de Souza et al., 2014). Recently, a *Croceicoccus* species has been found to be capable to produce AHL (Huang et al., 2015), which could be consistent with the slight increase in the AHL inducer genes during the rainy season. *Teredinibacter* have nitrogen fixation capabilities (Distel et al., 2002), while *Methanosalum* is a methylotrophic methanogen (Oren, 2014), which might aid in nutrient cycling during the rainy season. In contrast, the dry season core only contained the *Maledivibacter* genus, a member of the Clostridiales, Firmicutes. This genus produces hydrogen sulfide and ammonia under obligately halophilic conditions (Li, Zeng, et al., 2016). In fact, all the genera found in the seasonal cores are halophilic to some extent.

**Fig. 6:**
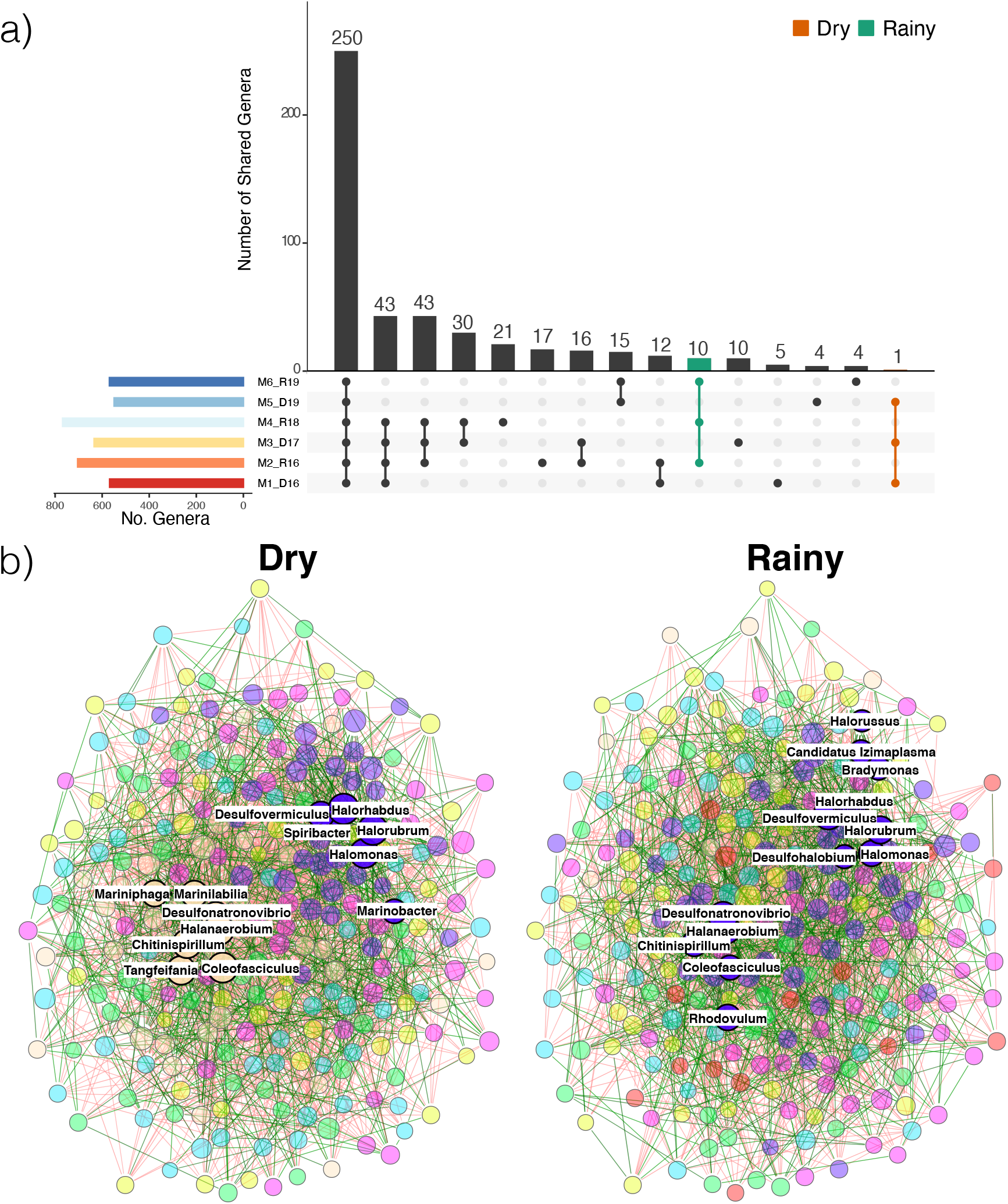
The core community at the Archean Domes microbial system. *a)* Upset plot showing the number of shared genera between different sample intersections. Global core showed 250 genera, while rainy core and dry core showed 10 and 1 genera, respectively. *b)* Co-occurrence networks for core genera during the dry and rainy seasons, where color indicates different clusters. Green and red edges represent positive and negative relationships, respectively. Hub taxa are shown with labels

We further analyzed the taxonomical structure and functions of the global core community. The core community consists of 250 genera, where most of them belong to the Proteobacteria (102), Bac-teroidetes (43), Firmicutes (28), Euryarchaeota (12), Actinobacteria (11), and Cyanobacteria (10) (Fig. S11: Supplementary material). Although these genera appear in every sample, their relative abundances fluctuate drastically between samples (Fig. S12: Supplementary material). For instance, *Coleofasciculus* transitioned from one of the genera with the highest abundance (10.2%) in 2017 to belong to the rare biosphere (0.06%) during the dry season of 2019. Most of the genera belonging to the core belong to the rare biosphere, although all the abundant genera belong to the core as well. Core functions relative abundances, on the other hand, appear highly conserved between samples (Fig. S11: Supplementary material), with processes such as carbohydrate, amino acid, protein, RNA, and DNA metabolisms being the most abundant ones. This result is consistent with the abundances of functions for the whole microbial system. PCoA ordination method suggest a seasonal pattern of functions, as the dry season samples of 2019 and 2016 were grouped in one cluster, while the rainy season samples of 2016 and 2018 were close to each other in another group. the dry season sample of 2017 and the rainy season sample from 2019 does not cluster to any of the aforementioned groups, and more samples will determine if these groups do preserve a seasonal pattern or not.

Core co-occurrence networks at the genus level also provide insights into the global core dynamics for both the dry and rainy seasons (Fig. 6). To begin with, there are 7 shared clusters of genera in both, dry and rainy seasons. Consistent with the networks built at the phylum level, the rainy season network for the 250 core genera displayed the addition of a new cluster that was not previously present in the dry season network. Additionally, many of the present genera relocate to different clusters between the dry and rainy seasons. This behavior could reflect how the core taxa differentially interact with each other in response to the environmental pressure. Even though these taxa are present in the whole community, regardless of the season, it is natural to infer that interactions within this core community are the ones that changes through the seasons. For both networks, global metrics were calculated, and once again, modularity and positive/negative ratio show consistency with the whole-community phylum networks (Table S7: Supplementary material); dry and rainy seasons displayed relatively high modularity values (0.17 and 0.22, respectively), and the slightly lower value during the dry season could reflect a drop in community stability during this state. Positive edges in both networks account for the ~41%, which result in high positive/negative ratios that further suggest a resistant and resilient community (Hernandez et al., 2021). Finally, hub taxa were predicted for each network, and among them, *Coleofasciculus*, *Chitinispirillum*, *Desulfonatronovibrio*, *Desulfovermiculus*, *Halanaerobium*, *Halomonas*, *Halorhabdus*, and *Halorubrum* are shared hubs between the seasons. Two hub groups appear during the dry season, whereas during the rainy season, every hub genus belong to the same cluster. It seems that some hub taxa (including *Desulfonatronovibrio*, *Coleofasciculus*, *Halanaerobium*, and *Chitinispirillum*) are also involved in the differential interactions between seasons. Given that sample size is small, detailed interaction analysis should be taken with caution. Thus, we retain ourselves to just an exploratory, non-conclusive, global analysis of these networks.

## 4 Discussion

Our metagenomic profiling from 2016 to 2019 at the Archean Domes aided us in the understanding of how microbial communities at this site are able to cope to environmental pressures during the dry and rainy seasons. Extended drought during roughly 9 months each year could be classified as a press disturbance, whereas daily temperature shifts amidst the desert could be interpreted as a pulse perturbation. Hence, the Archean Domes could be regarded as a multi-perturbation system.

Ecological resistance has been classically associated with the species/genera richness, where an increase or decrease in biodiversity could point to a decrease in compositional stability (Pennekamp et al., 2018). Nonetheless, there were none statistically significant differences for alpha diversity between the communities of the dry and rainy seasons. Moreover, we could infer that the community’s composition is heavily affected by each seasonal shift, pointing towards a sensitive, non-resistant, microbial community (Baho et al., 2012). Although community composition is constantly subject to taxonomic replacement, functions are mostly conserved, even though adjacent samples have shown more similarity in terms of function. Since functions are mostly preserved throughout the seasons, the highly diverse community might harbor a high degree of functionally redundant taxa (Allison and Martiny, 2008). This grants the community a robust capability to withstand taxonomic replacement, the product of a non-resistant community in terms of composition. Co-occurrence networks for the global (phylum level) and core (genus level) microbial networks show consistent results, as modularity and positive/negative ratio points towards a functionally stable community; in this sense, modules in these networks might reliably represent functional guilds or niche overlapping (Röttjers and Faust, 2018; Hernandez et al., 2021).

On the other hand, ecological resilience is not straightforward to assess in this system, as press disturbances are continuous and seasonal. Rather, inferences on ecological resilience could be evaluated under the assumption of many stable states in which a community may thrive. PCoA plots could be visualized as stability landscapes, where each snapshot of the community’s composition/function could be envisioned as a ball and where the different alternative stable states represent basins in the landscape. If resistance and resilience is high, a disturbance would not modify the current stable state of the community. On the contrary, if overall stability is low, and the disturbance powerful enough, the community will leave its current stable state to fall into an alternative stable state (Botton et al., 2006; Shade et al., 2012). PCoA plot for taxonomic composition (Fig. S7: Supplementary material), could be interpreted as the community transitioning towards different compositional states from 2016 until 2018, but in the 2019 samples the community remained in the same compositional state. Looking at the PCoA plot for functions (Fig. 5) we could see that seasonal “valleys” of alternative equilibrium are formed, whereas PCoA for the core functions, the community do return to the same seasonal stable state despite the press disturbance.

Community stability in this site has different components that contribute to the whole microbial resistance and resilience, at least functionally speaking. First, at the individual level, we found abundant genes related to salt and alkaline stress response which could confer physiological plasticity to the global core community, as these genera thrived despite the seasonal conditions. Mixotrophy is well represented in microbial communities and could further explain the individual functional plasticity between seasons (Eiler, 2006). At the population level, disturbances are well known to foster diversification, as these perturbations bring the community under selection pressures (Rainey and Travisano, 1998; Galand et al., 2016). The high diversity found in this pond might be the outcome of the multiple selection pressures acting on the microbial populations, providing an evolutionary adaptation; diversity overall at the community level promote functional resistance, since functional redundancy is expected in a genetically diverse community (Shade et al., 2012). For example, the Archean Domes microbial mats possess several carbon and energy metabolisms which could enhance the robustness of primary production and nutrient cycling. Finally, differential growth rates among the community members influence the overall composition, yet, we have seen that the microbial rare biosphere might play a key role in driving the community to new stable compositional states. Indeed, the rare biosphere drastically influence the taxonomic replacement in other systems (Jousset et al., 2017; Pascoal et al., 2021).

As stated previously, the Archean Domes microbial system is constantly subject to environmental stress, which could be arguably compared to those experienced by the Shark Bay microbial mats, a recognized terrestrial analog (Wong, White, et al., 2018; Campbell et al., 2020). Therefore, the Archean Domes site is potentially a promising analog for early Earth, and (plausibly) early Mars, just as other potential analogs found in the Cuatro Ciénegas Basin (López-Lozano et al., 2012; Moreno-Letelier et al., 2012; Souza et al., 2012). That being said, what can we learn of early life from these extant microbial communities?

Local and planetary disturbances throughout Earth’s history can be interpreted as environmental pulses or presses that perturbed the biosphere. Within this framework, microbial mats were biological structures that survived the onset of these threatening phenomena. Communities were probably compositionally sensitive, as shown by the microbial mats in this study and other microbial communities studied elsewhere (Allison and Martiny, 2008; Shade et al., 2012). While lineage extinction is expected as an effect of disturbances (Konhauser et al., 2015; Hodgskiss et al., 2019), multiple diversification events have been associated to post-disturbance episodes, such as genetic innovations driven by ocean-atmosphere oxygenation and changes in ocean chemistry (David and Alm, 2010; Chen, Sun, et al., 2020), or new clade emergence after major glaciation events (Chumakov, 2010). Therefore, Low compositionally resistant communities coupled with high mutation rates and further diversification within microbial mats could greatly influence the ceaseless search for alternative stable states in the stability landscapes. As major disturbances could completely modify the stability landscape (Shade et al., 2012), original compositional states could be never reached again, once the community is disturbed. Hence, it is plausible that modern composition of microbial mats (at least at the genus level) is substantially different from those during the Archean or Proterozoic Earth, and records of past compositional states might be unachievable.

As opposed to the continuous taxonomic replacement experienced in these systems, functional capability must have been a conservative feature for microbial mats throughout geologic time, as both resistance and resilience in functions were found for this analog site. Indeed, the fossil record show that modern metabolic capabilities could be traced back to past microbial mats and stromatolites (Buick, 1992; Bosak et al., 2009; Schopf, 2011; Lepot, 2020). This functional processes might be highly conserved across the community’s core, where functional redundancy is expected. Most modern microbial mat development is highly reliant on phototrophic cyanobacteria (Noffke, 2010; Prieto-Barajas et al., 2018), which lead us to wonder if microbial communities behaved similarly prior to the emergence of oxygenic photosynthesis. In this study, we find functional guilds that are stable to environmental perturbations, in which oxygenic photosynthesis is part of a local electron transfer circuit that includes energy and carbon metabolisms (Jelen et al., 2016). Closed electron transport circuits existed prior to the emergence of oxygenic photosynthesis, as depicted in Moore et al. (2017), where aerobic metabolisms emerged at a later stage in biological evolution. In this sense, microbial mats without oxygenic photosynthesis could rely on other metabolic processes to cycle nutrients and energy and become highly stable structures to functional changes. Future studies on modern microbial mats in hydrothermal vents (Rassa et al., 2009; Miranda et al., 2016) and phototrophic, anoxygenic sites (Visscher et al., 2020) would provide insights into this hypothesis.

Reconstructing past microbial ecologies, including their ecological stability, might provide valuable insights into the coevolution of the biosphere-geosphere. This knowledge has the potential to be applied to forecast microbial response under contemporary disturbances of global climate change, (Reinold et al., 2019), as well as potential modelling for microbial systems beyond Earth’s limits, where perturbations might be even more harsh than those experienced by terrestrial life.

## Supporting information

Functions groups kmeans

Functions raw counts

Taxonomy groups HDBSCAN

Taxonomy raw counts

## Competing interests

The authors declare no competing financial interests.

