## Supplementary material for "Modern microbial mats from the Chihuahuan Desert provide insights into ecological stability throughout Earth’s history"

#### Supplementary figures

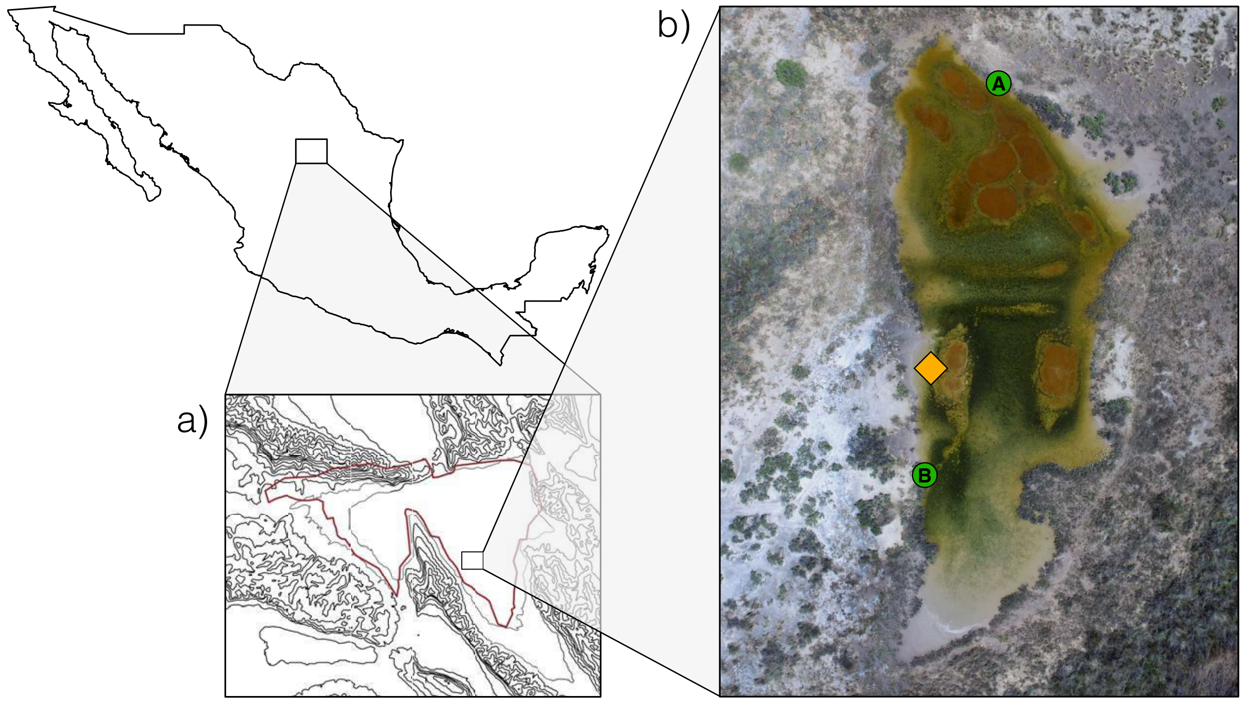


Fig. S1: Overview of the Archean Domes sample site. *a)* the Cuatro Ciénegas Basin, located in the Chihuahuan Desert. Depicted as a rectangle, the Pozas Azules ranch where the Archean Domes is located. *b)* Aerial view of the Archean Domes pond during the september 2019 sampling. The yellow diamond show the sampling point for the six metagenomes studied in this work. Green circles, A and B, indicate the picture location from Figure 1a and 1b, respectively.

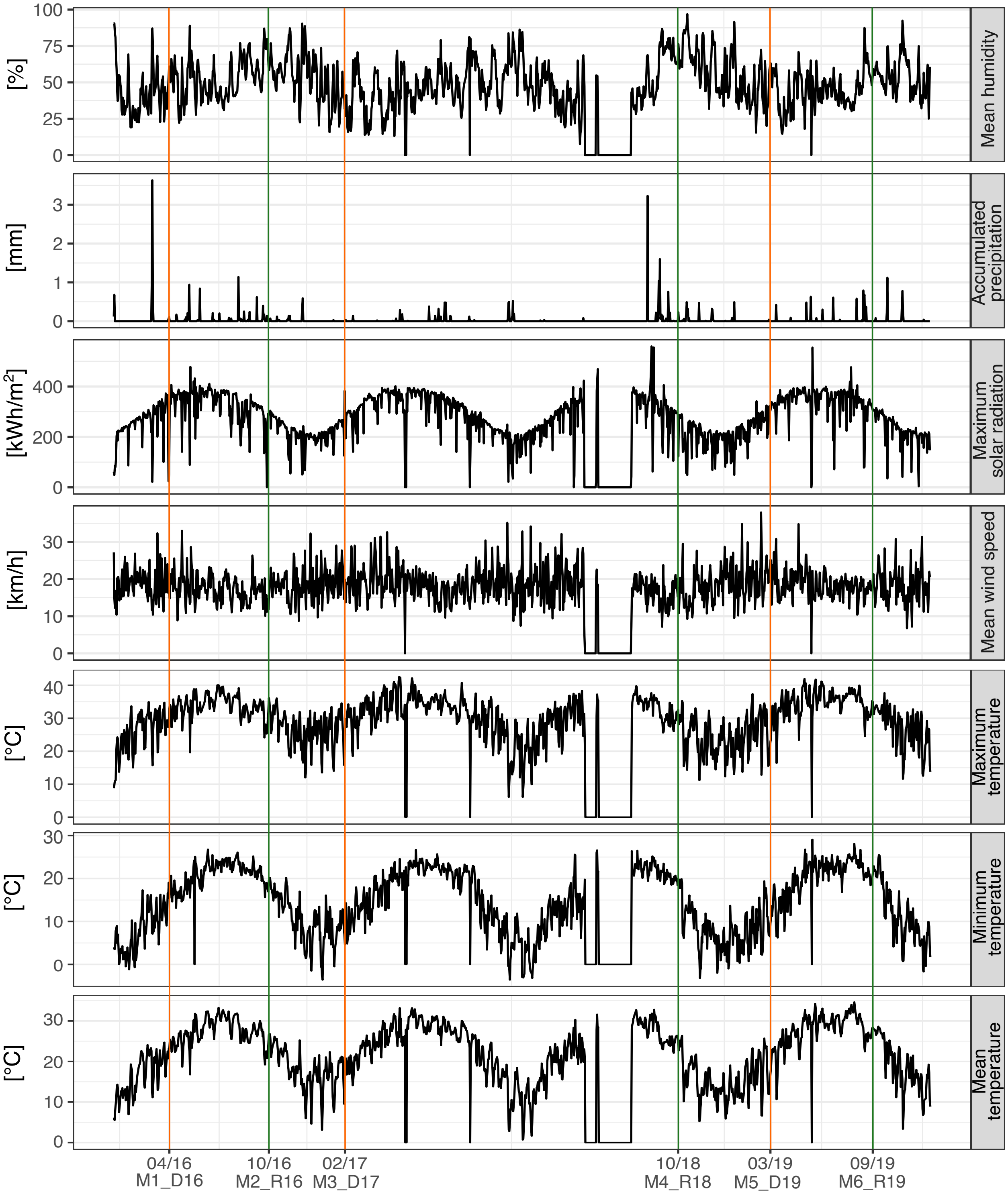


Fig. S2: Weather data retrieved from the EMA weather station No. 15DBB372, Cuatro Ciénegas, from 2016 to 2019. Data gaps during some months of 2018 represent that service was unavailable at that time. Lines show the sample day variables for each sample. Colors indicate rainy (green) and dry (orange) seasons.

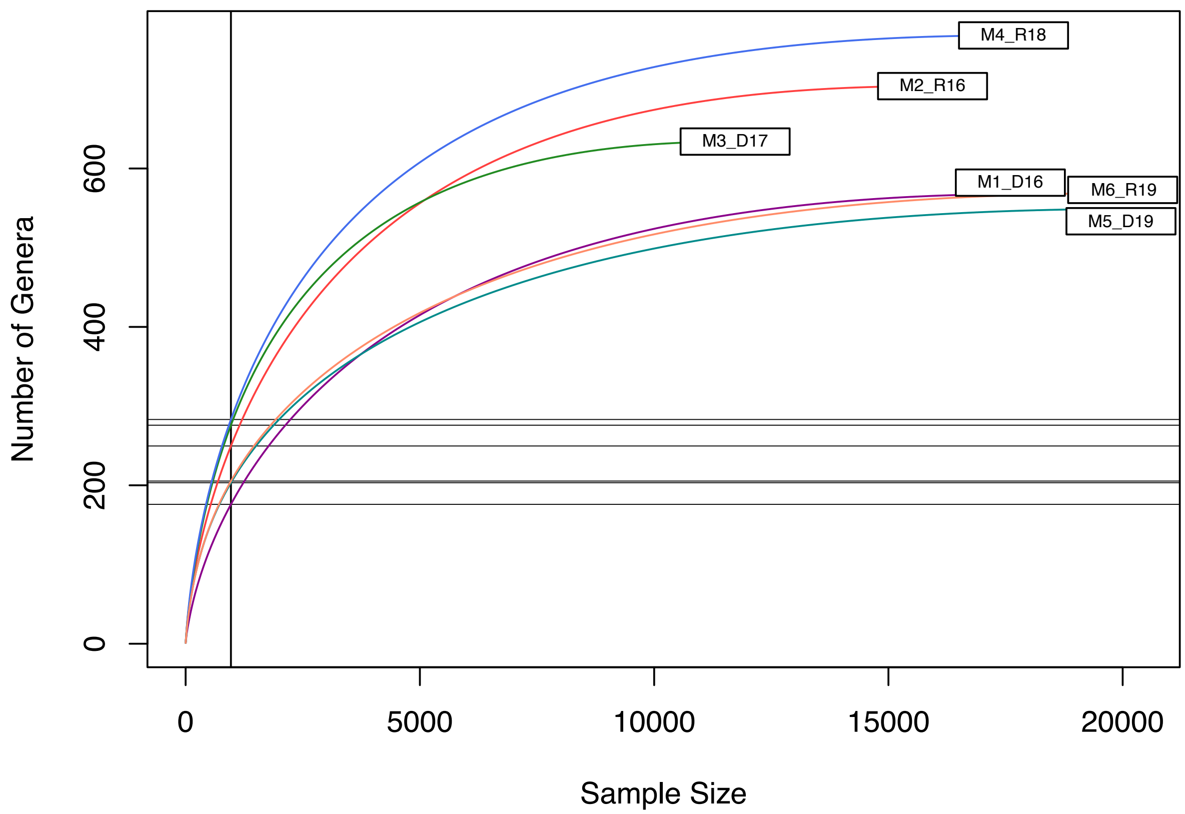


Fig. S3: Rarefaction curves for each sample studied. Sample size and number of genera are depicted in the horizontal and vertical axes, respectively. Each sample reaches saturation of genera richness and is suitable for sample comparison.

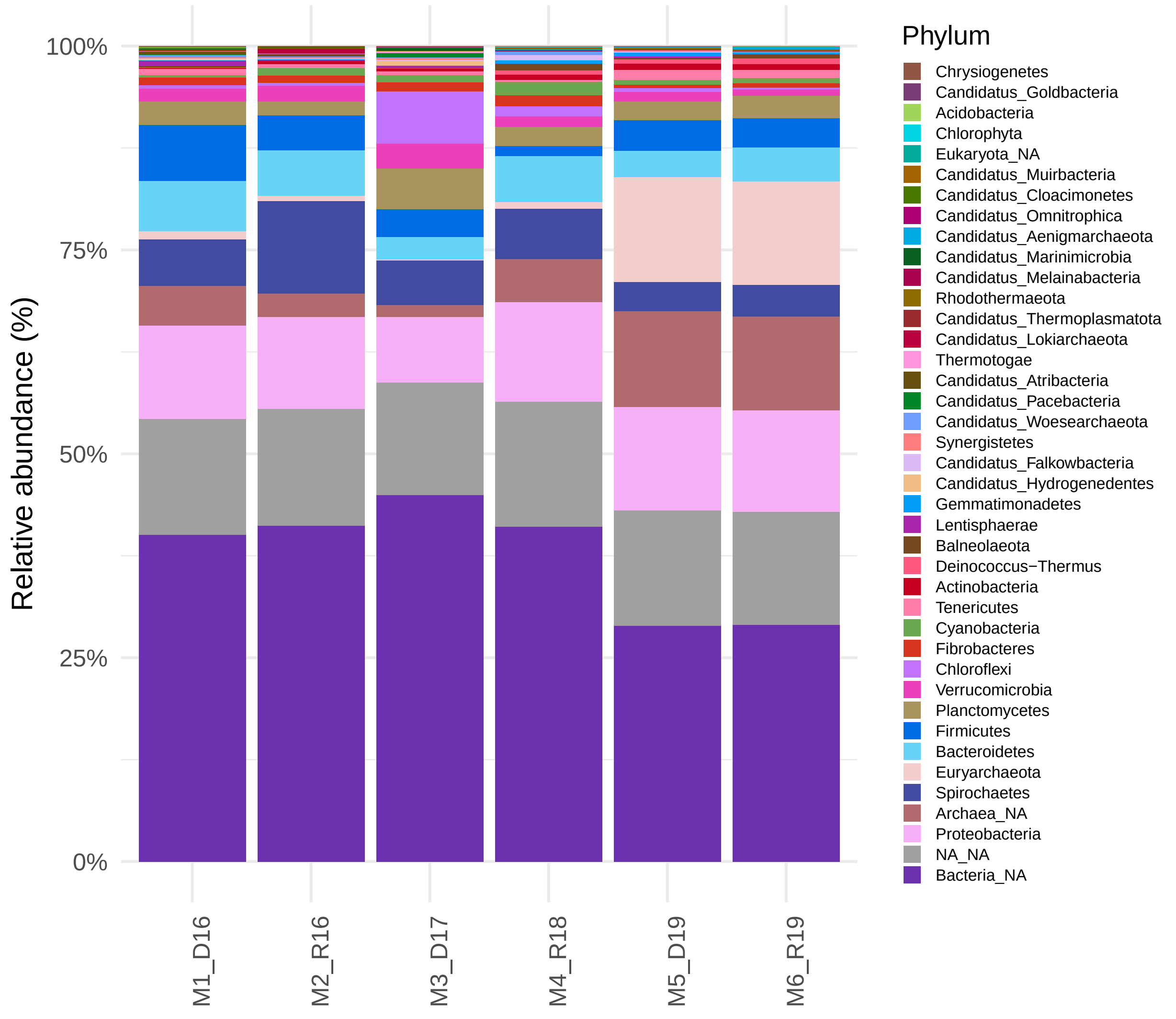


Fig. S4: Ribosomal-protein taxonomic profile of the Archean Domes system. Not annotated sequences were grouped in the NA categories.

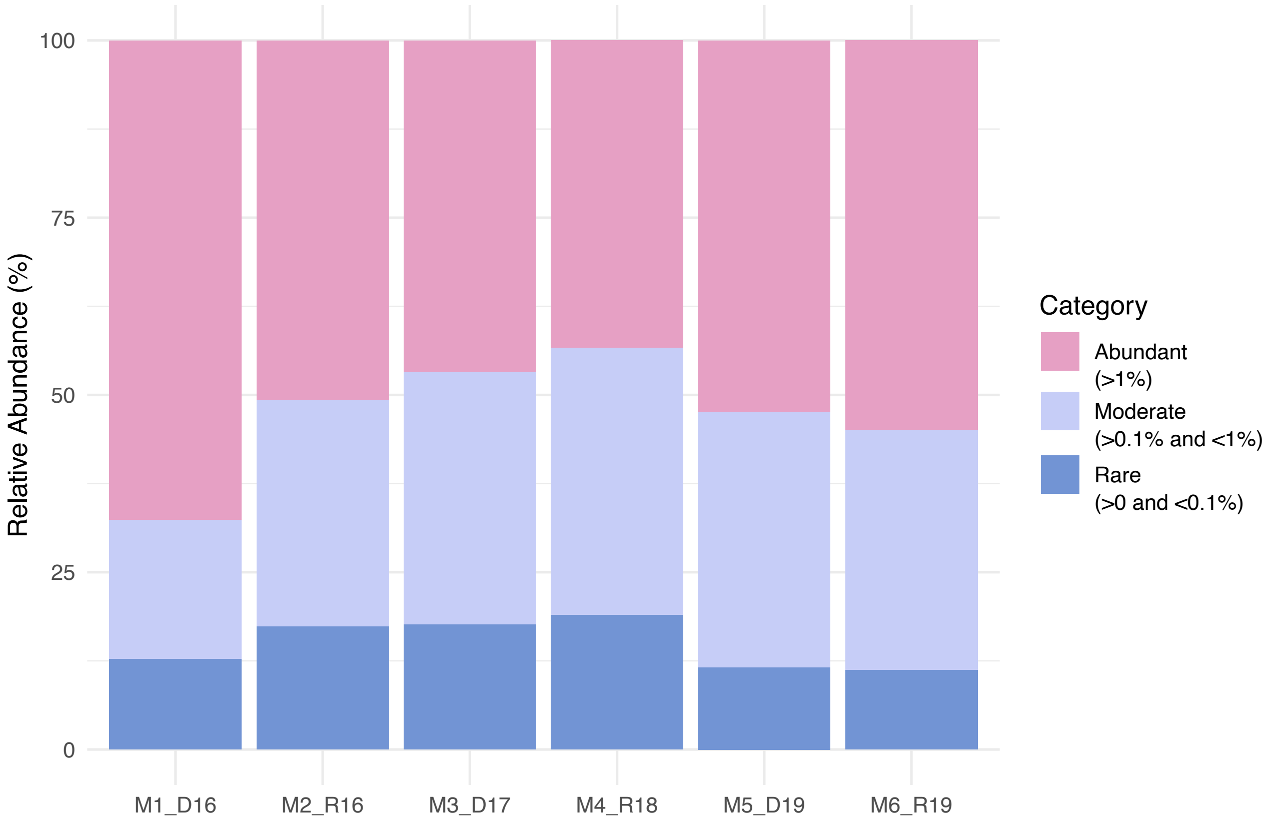


Fig. S5: The Archean Domes community based on their relative abundance at genus level. Abundant taxa comprise most of the whole community, in contrast to moderate and rare taxa abundance.

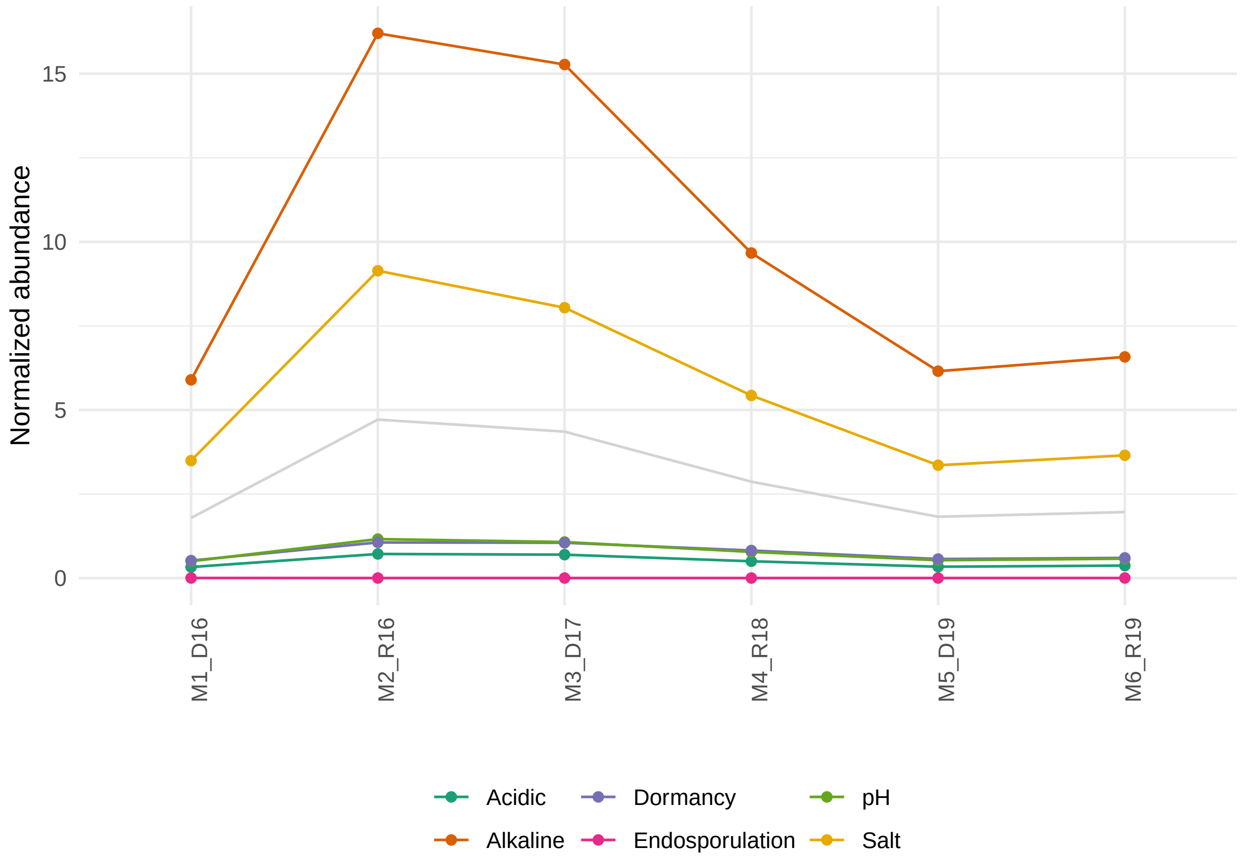


Fig. S6: Resistance genes found at the Archean Domes. Through all samples, most resistance genes associate to alkaline and salt response genes.

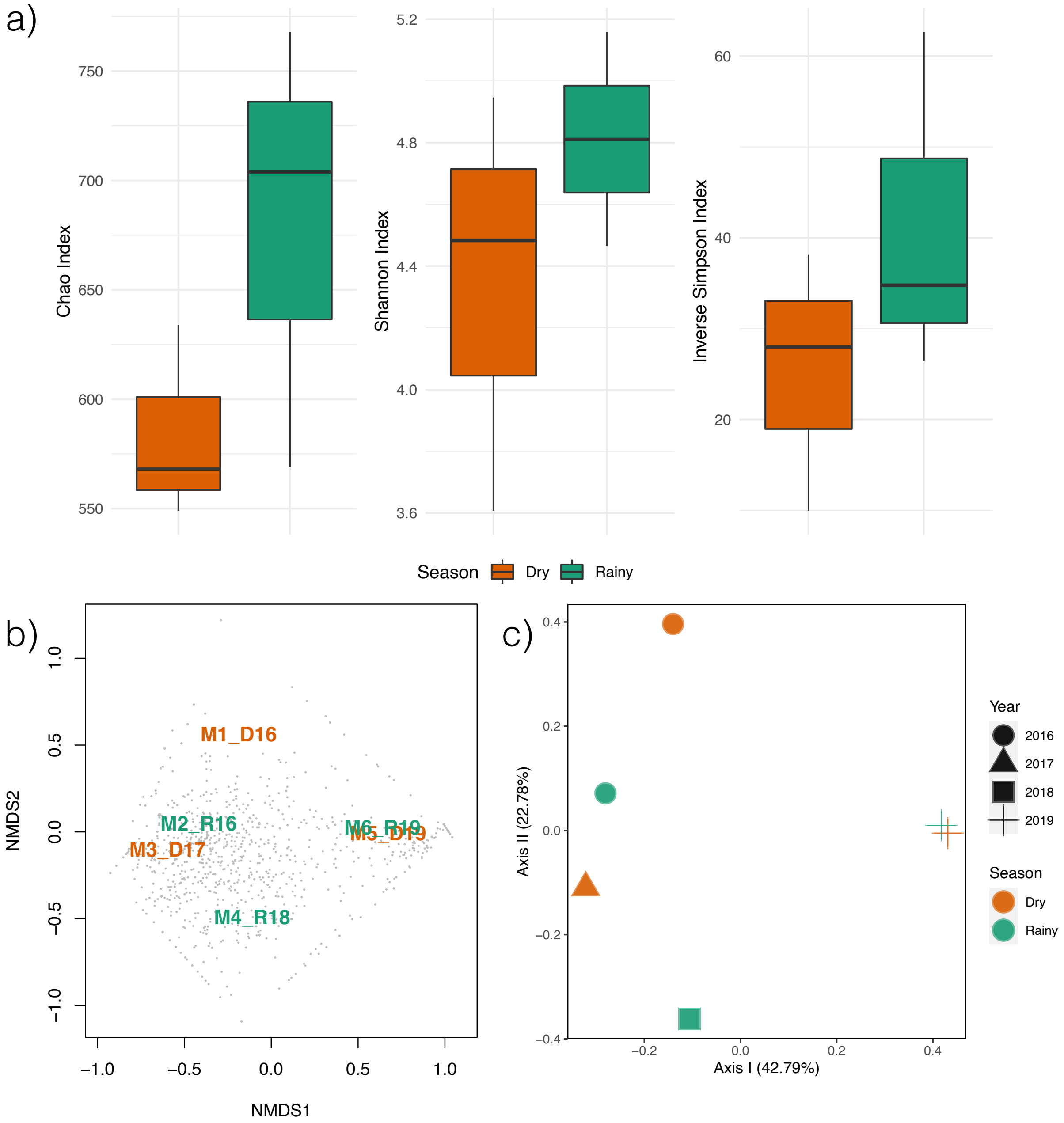


Fig. S7: Seasonal comparison of communities in the Archean Domes. *a)* Boxplots for diversity comparison for each seasonal state. performing Wilcoxon Rank Sum test showed no statistical differences between seasonal diversity. *b)* NMDS and *c)* PCoA analyses at genus level with Bray-Curtis measure. Both analyses showed no seasonal aggregation. NMDS and PCoA analyses at order level showed similar results to the ones at genus level.

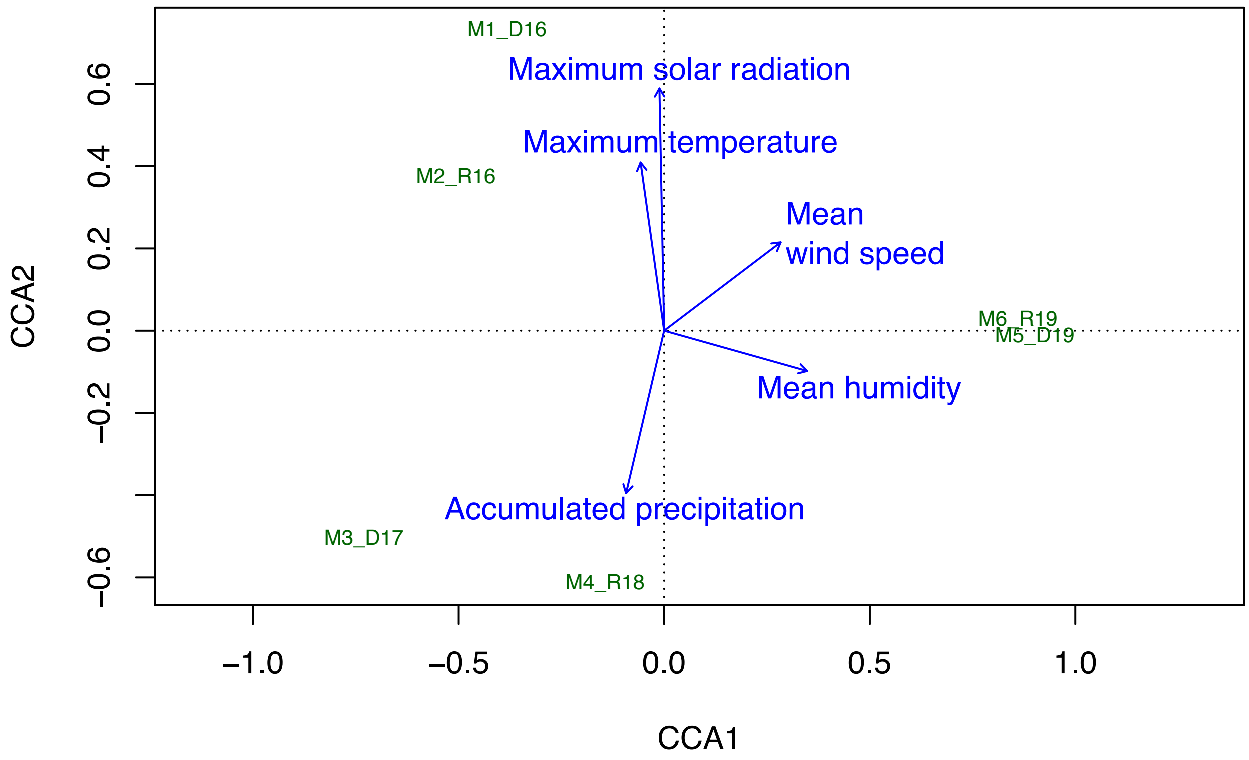


Fig. S8: CCA analysis with the environmental data provided by the EMA meteorlogical station. Meteorlogical data mean values for each sample month was input.

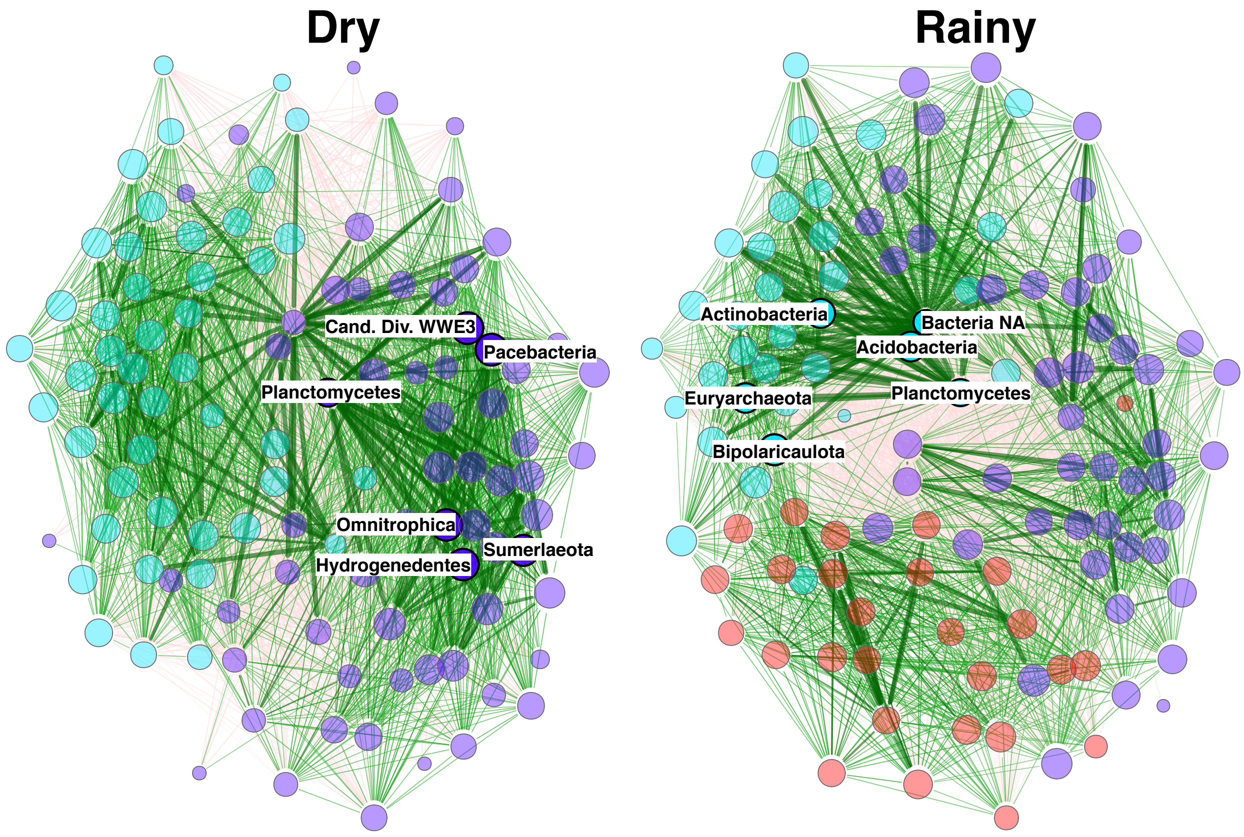


Fig. S9: Global network of shared phyla between seasons. Networks were built with the top 120 phyla across samples. Colors show clusters, and the same colors in both networks correspond to the same structural cluster. It can be appreciated the addition of a new cluster (red) during the rainy season. Green edges represent positive relationships, while red edges represent negative ones.

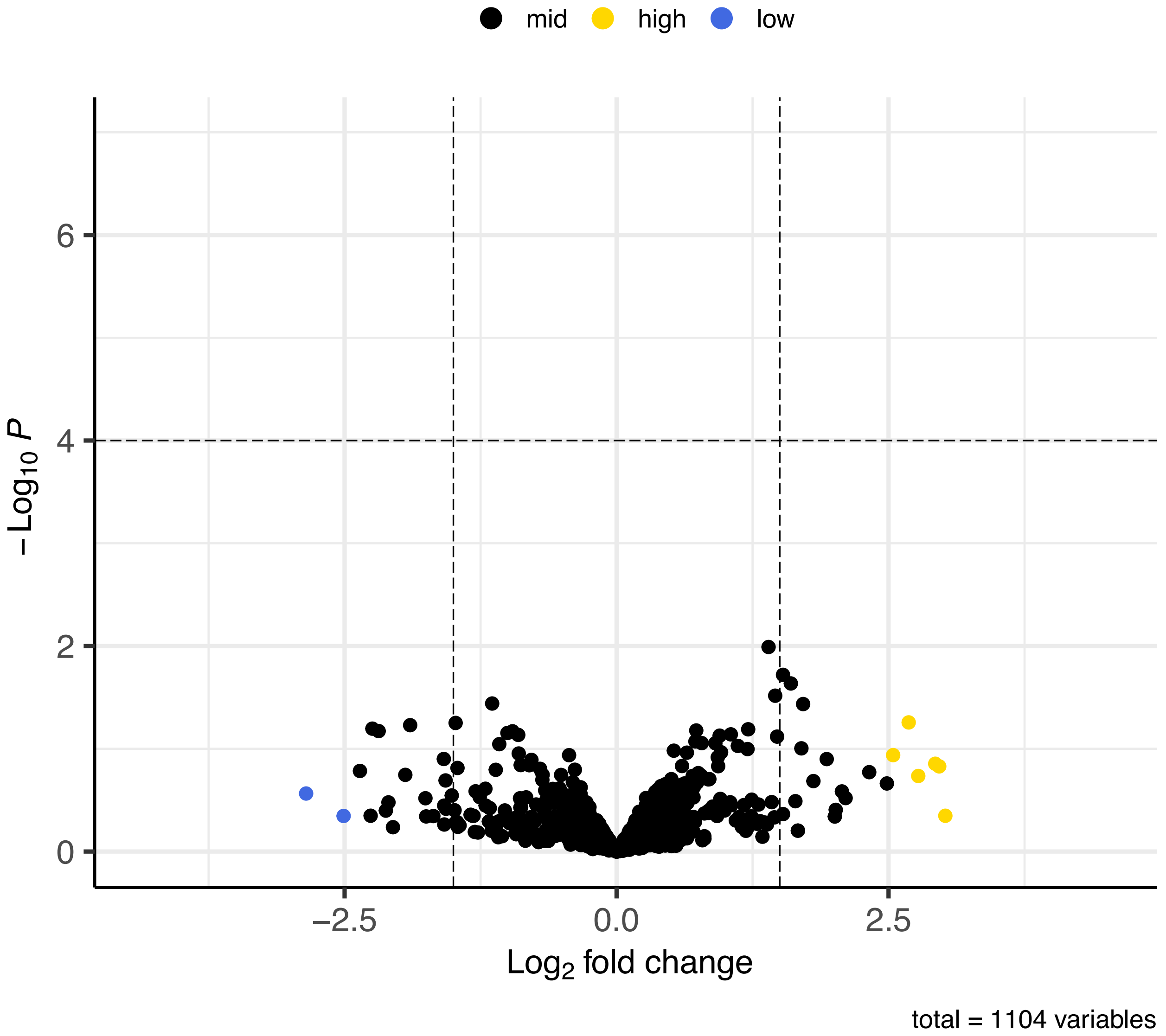


Fig. S10: Differential expression analysis of fuctions adapted to a metagenomic study (gene functions). No statistical significance was found for each function, although there are slight increases in both rainy (high) and dry (low) seasons.

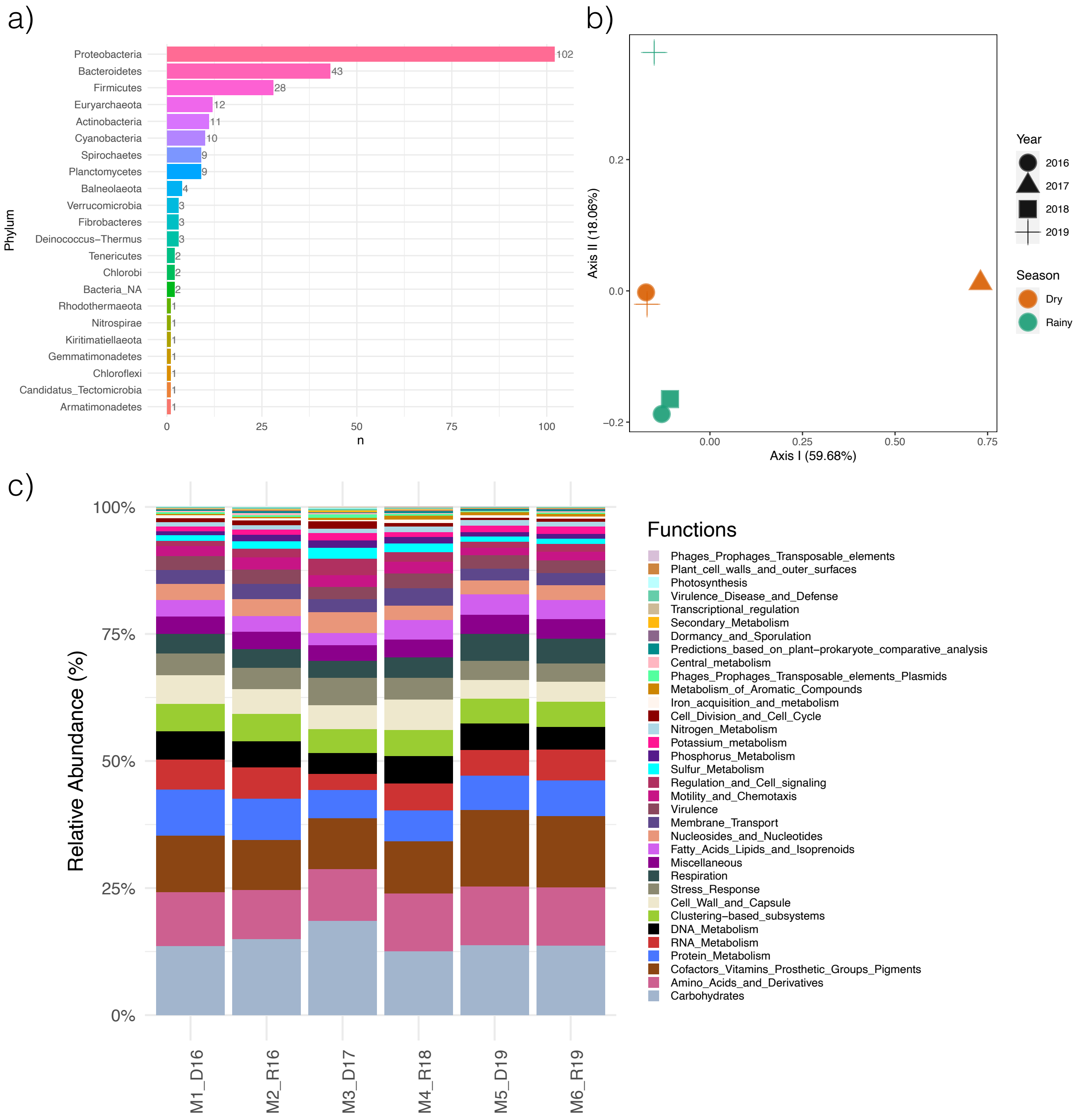


Fig. S11: Core taxonomic and functional details. *a)* Core composition at phylum level. *b* Core functions under a PCoA ordination method. Roughly, there appears to be a seasonal pattern with a close association between two dry and rainy season samples, although the rainy sample from 2019 and the dry sample from 2017 does not group with any other sample. *c)* Function composition for core genera. Overall, similar compositions between samples can be appreciated.

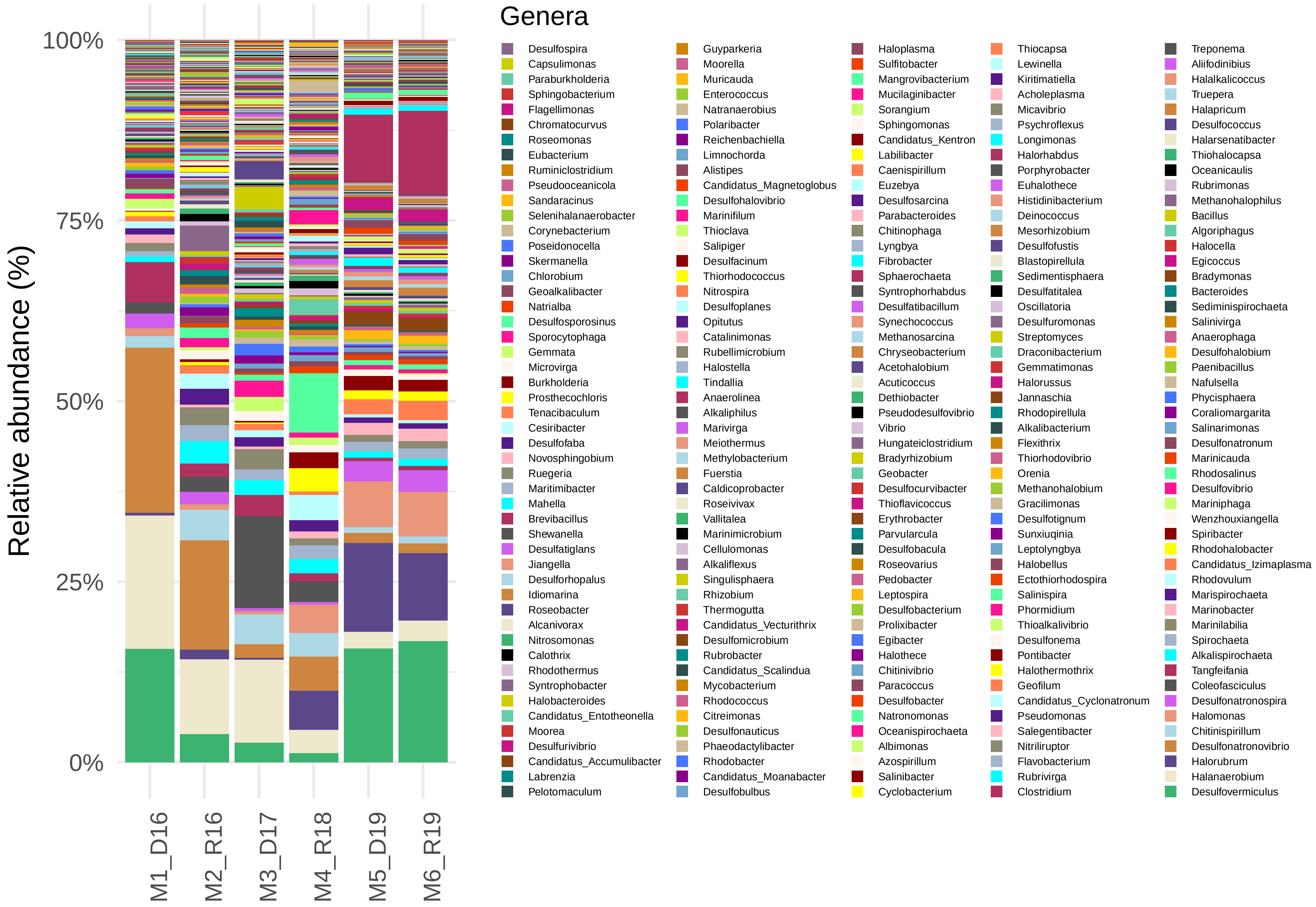


Fig. S12: Core composition at the genus level. Each of the 250 genera that constitute the core is shown in order to emphasize in the taxonomic replacement.

### Supplementary tables

Table S1: Sample-day weather parameters from EMA weather station No. 15DBB372, Cuatro Ciénegas.

| Sample/parameter | M1_D16 | M2_R16 | M3_D17 | M4_R18 | M5_D19 | M6_R19 |
|---|---|---|---|---|---|---|
| Sample day/time | 2016/04/09 9:48:38 | 2016/10/04 12:02:35 | 2017/02/17 11:38:28 | 2018/10/06 9:10:20 | 2019/03/20 11:32:19 | 2019/09/19 12:53:19 |
| Mean wind direction (°) | 153.5 | 149.5 | 245 | 170.5 | 123.5 | 111.5 |
| Mean wind speed (km/h) | 5.55 | 5.1 | 15.45 | 14.15 | 23.9 | 36.25 |
| Max. wind speed (km/h) | 14.3 | 12.1 | 43 | 22.5 | 39.1 | 63.2 |
| Min. Temperature (°C) | 23.1 | 28.1 | 21.1 | 24.6 | 16.8 | 29.5 |
| Mean Temperature (°C) | 23.9 | 29.2 | 22.53 | 25.13 | 17.3 | 29.73 |
| Max. Temperature (°C) | 24.8 | 29.8 | 23.1 | 25.9 | 17.9 | 29.9 |
| Mean humidity (%) | 63.83 | 50.5 | 28.5 | 70 | 45.67 | 54.15 |
| Mean pressure (mbar) | 923.52 | 925.07 | 927.82 | 928.43 | 935.37 | 929.6 |
| Max. solar radiation (kWh/$m^2$) | 721 | 859 | 864 | 607 | 971 | 942 |

Table S2: Raw reads and quality control (QC) metadata for each shotgun metagenome studied.

| Sample/ parameter | Raw reads | Low QC reads | Total reads | Paired reads | Forward Unpaired | Reverse Unpaired |
|---|---|---|---|---|---|---|
| M1_D16 | 28,859,454 | 2,425,863 | 26,433,591 | 21,012,271 | 4,755,240 | 666,080 |
| M2_R16 | 4,772,053 | 287,632 | 4,484,421 | 3,722,418 | 671,964 | 90,039 |
| M3_D17 | 8,203,484 | 790,818 | 7,412,666 | 5,915,153 | 1,255,747 | 241,766 |
| M4_R18 | 10,030,782 | 1,561,639 | 8,469,143 | 6,115,153 | 2,098,789 | 255,201 |
| M5_D19 | 25,873,990 | 3,589,352 | 22,284,638 | 17,676,424 | 3,645,559 | 962,655 |
| M6_R19 | 20,153,088 | 1,471,902 | 18,681,186 | 16,378,340 | 1,784,198 | 518,648 |

Table S3: Metagenome assembly metadata for each sample. Minimum, average, and maximum contig length is also shown

| Sample/ parameter | Total base pairs | Assembled (bp) | Not assembled (bp) | Min. contig length (bp) | Avg. contig length (bp) | Max. contig length (bp) |
|---|---|---|---|---|---|---|
| M1_D16 | 9,838,571,624 | 857,058,612 | 8,981,513,012 | 500 | 1,274.80 | 159,881 |
| M2_R16 | 1,491,700,706 | 190,580,164 | 1,301,120,542 | 500 | 1,243.70 | 86,189 |
| M3_D17 | 2,627,434,540 | 304,829,428 | 2,322,605,112 | 500 | 1,291.70 | 121,677 |
| M4_R18 | 2,985,543,837 | 329,842,768 | 2,655,701,069 | 500 | 1,089.90 | 75,330 |
| M5_D19 | 7,799,883,462 | 588,197,199 | 7,211,686,263 | 500 | 1,142.20 | 142,626 |
| M6_R19 | 7,744,171,739 | 621,571,725 | 7,122,600,014 | 500 | 1,108.90 | 124,235 |

Table S4: Metagenome assembly metadata for sequences not initially assembled.

| Sample/ parameter | Assembled sequences | Not assembled sequences (Forward) | Not assembled sequences (Reverse) |
|---|---|---|---|
| M1_D16 | 672,295 | 1,263,688 | 197,513 |
| M2_R16 | 153,232 | 244,559 | 26,150 |
| M3_D17 | 235,985 | 323,183 | 66,130 |
| M4_R18 | 302,633 | 735,840 | 91,703 |
| M5_D19 | 515,141 | 888,820 | 242,019 |
| M6_R19 | 560,549 | 427,091 | 121,222 |

Table S5: Taxonomic annotation metadata for each sample and relative abundances for each domain of life.

| | Reads processed | | | | Relative abundance (%) | | | |
|---|---|---|---|---|---|---|---|---|
| Sample/parameter | Total Found | Unclassified | Classified | Phylotypes | Archaea | Bacteria | Eukaryota | Viruses |
| M1_D16 | 1,562,120 | 1,083,248 | 478,872 | 8,699 | 3.06 | 96.65 | 0.20 | 0.08 |
| M2_R16 | 1,288,875 | 578,442 | 710,433 | 10,969 | 4.27 | 95.45 | 0.17 | 0.11 |
| M3_D17 | 285,524 | 190,940 | 94,584 | 3,872 | 1.44 | 98.38 | 0.13 | 0.05 |
| M4_R18 | 1,791,166 | 867,900 | 923,266 | 12,370 | 7.27 | 92.01 | 0.51 | 0.21 |
| M5_D19 | 1,833,943 | 1,020,352 | 813,591 | 10,079 | 33.61 | 65.83 | 0.19 | 0.36 |
| M6_R19 | 2,142,378 | 884,359 | 1,258,019 | 11,840 | 33.60 | 65.83 | 0.19 | 0.39 |

Table S6: Alpha diversity indexes for each sample studied.

| Sample/Index | Chao | Shannon | Simpson | Inverse Simpson |
|---|---|---|---|---|
| M1_D16 | 271.2500 | 2.792135 | 0.8184993 | 5.509622 |
| M2_R16 | 143.4000 | 2.642506 | 0.8068319 | 5.176839 |
| M3_D17 | 200.0833 | 2.592936 | 0.7840716 | 4.631164 |
| M4_R18 | 205.1154 | 2.716131 | 0.8082304 | 5.214591 |
| M5_D19 | 233.0000 | 3.128192 | 0.8797076 | 8.313074 |
| M6_R19 | 231.3750 | 3.077123 | 0.8787593 | 8.248053 |

Table S7: Network metrics for global network (phylum level) and core network (genus level). Hub taxa found by NetCoMi is also shown.

| | Phylum-level Global network | | Genus-level Core network | |
|---|---|---|---|---|
| Category/ parameter | Dry season | Rainy season | Dry season | Rainy season |
| Number of components | 1.0 | 1.0 | 1.0 | 1.0 |
| Clustering coefficient | 0.75980 | 0.73311 | 0.11670 | 0.12789 |
| Modularity | 0.01279 | 0.06986 | 0.16715 | 0.21577 |
| Positive edge percentage | 49.01681 | 48.77283 | 41.94115 | 41.41117 |
| Edge density | 0.57415 | 0.56764 | 0.07316 | 0.07650 |
| Natural connectivity | 0.21904 | 0.20453 | 0.01452 | 0.01545 |
| Vertex connectivity | 3.0 | 1.0 | 6.0 | 7.0 |
| Edge connectivity | 3.0 | 1.0 | 6.0 | 7.0 |
| Average dissimilarity | 0.78332 | 0.78216 | 0.97908 | 0.97812 |
| Average path length | 0.82592 | 0.84478 | 1.45398 | 1.44128 |
| Hub taxa | cand.div. WWE3<br>Hydrogenedentes<br>Omnitrophica<br>Pacebacteria<br>Planctomycetes<br>Sumerlaeota | Acidobacteria<br>Actinobacteria<br>Bacteria NA<br>Bipolaricaulota<br>Euryarchaeota<br>Planctomycetes | Chitinispirillum<br>Coleofasciculus<br>Desulfonatronovibrio<br>Desulfovermiculus<br>Halanaerobium<br>Halomonas<br>Halorhabdus<br>Halorubrum<br>Marinilabilia<br>Mariniphaga<br>Marinobacter<br>Spiribacter<br>Tangfeifania | Bradymonas<br>cand. Izimaplasma<br>Chitinispirillum<br>Coleofasciculus<br>Desulfohalobium<br>Desulfonatronovibrio<br>Desulfovermiculus<br>Hlanaerobium<br>Halomonas<br>Halorhabdus<br>Halorubrum<br>Halorussus<br>Rhodovulum |
